# Aphid Salivary MIF Modulates Plant Programmed Cell Death and DNA Damage Response and Interacts with SOG1

**DOI:** 10.64898/2026.04.01.715815

**Authors:** Killian Menuet, Carlotta Aurora Lupatelli, Ariane Fazari, Thierry Fricaux, Georges De Sousa, Janice de Almeida Engler, Christine Coustau

## Abstract

The establishment of aphid-plant interaction involves the secretion of a salivary MIF protein. Morphological analyses revealed that aphid MpMIF1 prevents plant cell death, protects organelles from stress, and may promote plant cellular recovery. Co-expression of aphid MpMIF1 and the cell death inducer Npp1 revealed that MpMIF1 modulates autophagy-related genes ATG7/BECLIN1, impair plant senescence regulator ATAF1 and regulate apoptosis-like via Caspase-3-like activity. This effect on multiple-cell death pathways helps to maintain cellular homeostasis during aphid infection. Investigations on DNA Damage Response (DDR) signaling pathways demonstrated that aphid MpMIF1 reduces γH2A.X phosphorylation, maintains activity of the DNA repair protein RAD51 and stabilizes cell cycle checkpoint expression WEE1 under genotoxic stress. Therefore, MpMIF1 actively participates to the maintenance of a functional DDR. Finally, we showed that aphid MpMIF1 physically interacts with SOG1, a functional analog of animal p53 and central regulator of DDR, cell cycle arrest and programmed cell death in plants. These findings establish MpMIF1 as a key regulator of plant cell death during aphid–plant interactions and highlight its potential as a biotechnological tool for protecting major crops against aphid infection.

## 1. Introduction

Every living organism, whether plant, animal, fungus, or microbe, serves as host for various parasitic species [1]. Among plant parasites, aphids, represent a global threat to agriculture due to their ability to drain nutrients from plants via the phloem and to transmit plant-pathogenic viruses [2]. Aphid’s feeding involves insertion of the stylet between epidermal cells, followed by probing of mesophyll cells, long-repeated probing of companion cells and their associated phloem cells, and finally, insertion in the phloem cells for phloem sap ingestion [3]. During these probing phases, aphids both uptake small samples of plant cell cytoplasm and secrete salivary molecules interacting with plant defense responses and cell death processes [3–5]. The mechanical damages associated with this feeding behavior is known to induce cellular alterations such as plasmolysis [6,7], thereby highlighting the important role of secreted salivary molecules in preserving cell viability of the feeding site. One of the key salivary proteins has been identified as a Macrophage Migration Inhibitory Factor (MIF) [8]. The MIF protein family comprises evolutionarily ancient proteins found across diverse biological kingdoms, including vertebrates, invertebrates, protists, fungi and plants [9–11]. In both vertebrates and invertebrates, MIFs play key roles in orchestrating the innate immune response and tissue repair, favoring cell proliferation and inhibiting cell death [12–14]. The secretion of MIF proteins had also previously been documented in parasites of vertebrate species, including nematodes (e.g., hookworms) [15,16], protozoa (e.g., *Leishmania* and *Plasmodium*) [17], and ticks [18] and shown to contribute to the modulation of the vertebrate host immune responses to facilitate infection [15,17].

Aphids possess a multigenic family of MIF proteins whose expression is differentially modulated during a beneficial or deleterious interaction (i.e. with mutualistic symbionts or parasites /pathogens) [19]. Interestingly, a single MIF member is expressed in the salivary glands, suggesting that one of these immune regulators has been rerouted to participate to the plant-aphid interaction [8]. These salivary MIFs have been shown to be secreted during feeding in both the pea aphid, *Acyrthosiphon pisum*, and the green peach aphid *Myzus persicae* [8]. Salivary secretion of both MIF proteins (ApMIF1 and MpMIF1) was shown to be crucial for infection of the plant host as their expression knock-down resulted in the failure of aphid feeding, establishment and reproduction [8]. Furthermore, the ectopic expression of aphid MpMIF1 in leaf tissues was shown to inhibit major plant immune responses, such as the expression of defense-related genes, callose deposition, and cell death [8,20]. These results represented the first report of an animal MIF modulating plant immune responses and cell death [8].

The intriguing question emerging from this ability of aphid MIF to modulate plant immune response and inhibit plant cell death is the question of the molecular processes involved. In animals, it is well established that MIF can suppress the p53-dependent apoptotic pathway [21] by preventing p53 translocation from the cytoplasm to the nucleus through direct intracellular interaction [22–24]. Additionally, MIF can activate the ERK1/2 MAP kinase (MAPK) signaling pathway via specific binding to the extracellular domain of CD74, along with the recruitment of the CD44 co-receptor, promoting cell proliferation and inhibiting programmed cell death (PCD) [25,26]. However, these major partner molecules such as CD74 or p53 are missing in plants. The aim of this study was therefore to explore the effect of aphid MpMIF1 on plant cell death using complementary approaches targeting the known plant PCD pathways, and the known and conserved MIF pathways.

In plants, PCD plays a critical role in numerous developmental processes, including gametophyte maturation, xylem treachery element differentiation and root development. PCD is also a key component of plant immune responses, particularly during the hypersensitive response (HR) triggered by pathogen attack [27,28]. In contrast to animal PCD processes and molecular pathways that are precisely described and classified, plant PCD processes are less clearly characterized leading to a complex and often ambiguous categorization [29,30]. By integrating current morphological and molecular criteria, and by analogy with the animal PCD processes, plant PCD can be broadly categorized into three types: apoptosis-like cell death (Type I), autophagic-like cell death (Type II), and necrosis-like (Type III) [28,31]. Other forms of plant cell death, such as ferroptosis-like, vacuolar cell death and hypersensitive response, can generally be classified within these three overarching categories based on their molecular and morphological features. For instance, vacuolar cell death involves autophagic processes and enzymes with caspase-like activity, such as vacuolar processing enzymes (VPEs) and metacaspases [32,33]. Similarly, the hypersensitive response (HR) shares morphological and molecular characteristics of animal apoptosis [34], pyroptosis and necroptosis [35,36], with involvement of evolutionarily conserved AuTophaGy-related (ATG) proteins such as ATG7 [37] and BECLIN1/ATG6 [38].

Recently, the concept of apoptosis cell death in plants has been increasingly challenged. Plant cell structure prevents their fragmentation and the formation of apoptotic bodies [35,39,40]. However, caspase-like proteases have been identified as key participants in plant PCD, and numerous studies continue to refer to “apoptosis-like” processes in plants. This is due to the fact that plant PCD exhibits many of the hallmark biochemical and morphological features of apoptosis, including cell shrinkage and chromatin condensation [41]. Therefore, because this debate is still unresolved, we will use the term ‘apoptosis-like’ in this article. Among the various internal and external factors that can trigger PCD, DNA damage as Double-Strand Breaks (DSBs) is particularly harmful to the cell [42]. In both plants and animals, the DNA Damage Response (DDR) pathways are highly conserved, although key differences exist [43]. Members of the phosphoinositide 3-kinase-related kinase (PIKK) family, such as Ataxia Telangiectasia Mutated (ATM) and Rad3-related (ATR) are rapidly activated in response to DNA damage [44]. The MRN complex is responsible for recruiting ATM/ATR to DSBs sites. Upon recruitment, ATM phosphorylates the histone variant H2AX [45], leading to the formation of γH2AX. γH2AX serves as both a marker for DNA damage and a scaffold for the recruitment of additional DDR proteins [46], including key players such as RAD51 and BRCA1, which are essential for homologous recombination repair mechanisms [47,48]. One of the most significant differences between animals and plants in their DNA damage response lies in the identity of the primary downstream effector. In animals, p53 plays a central role in determining whether a cell undergoes cell-cycle arrest, initiates DNA repair, or proceeds to apoptosis following DNA damage. However, no homolog of p53 has been identified in any model plant species [49,50].

Curiously, SOG1 (SUPPRESSOR OF GAMMA RESPONSE 1) has been proposed as the functional analog of mammalian p53 in plants [51,52]. SOG1 and p53 share no sequence similarity, indicating that SOG1 likely evolved independently as a plant-specific regulator of DDR [51,52]. SOG1 belongs to the plant-specific NAC domain family of transcription factors, and it plays a crucial role in coordinating the transcriptional response to DNA damage, regulating PCD, and enforcing cell cycle checkpoints [42,52]. Notably, members of the NAC transcription factor family (NAM, ATAF1/2, and CUC2) are central to the regulation of senescence [53]. For example, ATAF1 functions as a positive regulator of senescence [54].

Therefore, in response to genotoxic stress, cell cycle progression is often delayed or arrested at key checkpoints [51,52]. WEE1 kinase is one of the key effectors involved in halting cell cycle progression following DNA damage, and its activation is mediated by ATR/ATM and SOG1 [55,56].

To better understand the role of the aphid MpMIF1 protein in inhibiting plant PCD and its potential involvement in DDR mechanisms, we performed *in planta* agroinfiltration using the Npp1 protein from *Phytophthora parasitica* as a tool to induce localized cell death in *Nicotiana benthamiana* leaves. It has been demonstrated that NLPs, including Npp1, function as cytolytic toxins that trigger plasma membrane disruption by binding to specific plant sphingolipid receptors [57,58]. Through the formation of pores in the membrane, they cause osmotic collapse of the plant cell, ultimately leading to cell death [58]. This generalized mode of action makes Npp1 a valuable tool for studying plant cell death broadly, without prior assumptions about specific pathways. In this study, we conducted a comprehensive set of morphological, molecular, and biochemical analyses to investigate the impact of MpMIF1 on plant PCD and DDR under cell death conditions induced by Npp1. We co-expressed Npp1 with MpMIF1 proteins to assess the potential effect of MpMIF1 on plant PCD and DDR markers, on plant MAPKs Erk1/2 pathways, and we investigated the potential binding of this insect-derived MIF with the plant functional analog of p53, that is SOG1.

Our findings highlight MpMIF1 as a key factor in inhibiting plant cell death during aphid–plant interactions, underscoring its potential as a valuable biotechnological tool for protecting important crops against this pest.

## 2. Materials and Methods

### 2.1. Plant material and expression constructs for transient Agrobacterium-mediated transformation

Seeds of *Nicotiana benthamiana* were grown in pots containing mixed sand and soil (1:5) under 16 h light/8 h darkness photoperiod at 21/23°C. Leaves from 5–6-week-old plants of *N. benthamiana* were syringe-infiltrated with *Agrobacterium tumefaciens* transformants and adapted as described below. Necrosis-inducing *Phytophthora protein 1* (*Npp1*, GenBank: XM_008912603.1) and *Myzus Persicae MIF1* (*MpMIF1* GenBank: KP218519.1) sequences were cloned into the entry vector pDON207 and transferred into the destination vectors pH2GW7.0 and pK7WGF2.0 vector using the Invitrogen BP and LR reaction protocols. The constructs were validated by sequencing. As control, Empty Vector of pH2GW7.0 and pK7WGF2.0 vectors already containing GFP sequence were used. DH5a competent Cells were transformed and selected by antibiotic resistance. Plasmids were then transformed by electroporation into *Agrobacterium tumefaciens* strain GV3101. Transformants were selected using gentamycin (25μg/mL), rifampicin (50μg/mL) and spectinomycin (100μg/mL). Recombinant strains were grown in LB medium with the above-mentioned antibiotics at 28°C to an OD600 of 0.4. After centrifugation, the pellet was resuspended in Agro-mix and incubated at room temperature (10 mM MgCl2, 10 mM MES pH 5.6, 150M Acetosyringone). Empty Vector was used as a control of the bacteria infiltration and MpMIF1+GFP to control the success of the infiltration and to check for a potential effect of two proteins express at the same time. The subcellular localization of MpMIF1 within plant cells following agroinfiltration is shown in Figure S1 and is consistent with the expected secretion of aphid MIF in plant cells and with the nucleocytoplasmic expression of MIF proteins reported in many species [11,59]. Npp1+GFP was used to investigate the effect of cell death inducer plus a second non-relevant protein and Npp1+MpMIF1 was used to investigate the effect of cell death inhibition by MpMIF1. All co-infiltrations were used at a 1:1 ratio.

### 2.2. Chlorophyll quantification

Infiltrated leaf discs were harvested at 1-, 2- and 3-days’ post infiltration (DPI) and deposited in a 96-well ELISA/RIA (Falcon®) plate containing 300 μL of absolute ethanol in each well. The plate was covered and placed at 4°C over-night and 16 hours later 200 μL were transferred to a new 96-well plate. The absorbance was subsequently measured for each sample at three different wavelengths: 750, 647 and 664 nm by a spectrophotometer (SpectraMax Plus 384).

### 2.3. LysoTracker staining

Leaf sections were stained with 5 µM LysoTracker™ Red DND-99 (1 mM stock in TRIS buffer, 50 mM, pH 6.8) for 10 minutes, then washed in TRIS for 20 minutes. Samples were mounted and imaged using a Zeiss LSM780 confocal microscope with a 20× Plan-Apo objective. Excitation was performed with 488 nm (argon) and 561 nm (DPSS) lasers. Emissions were collected in three channels: 493–555 nm (GFP), 568–631 nm (LysoTracker), and 655–711 nm (chlorophyll A), using two spectral detectors. Z-stacks and time series were acquired using ZEN Blue software and processed by maximum intensity projection. Vesicles were counted in ImageJ and normalized per cell. Experiments were repeated six times, and statistical analysis was performed using GraphPad Prism.

### 2.4. Propidium iodide staining

Infiltrated leaf samples were harvested, observed by a ZEISS AxioZoom.v16 fluorescence macroscope with PlanNeoFluar Z 1.0x/0.25 objective under bright-field illumination. To visualize dead cells with fluorescent nuclei, samples were stained with Propidium Iodide (PI). Fresh tissue was incubated in a PI solution (1 µg/mL in distilled water with 0.02% DMSO) for 1 hour in the dark. After staining, samples were mounted in ∼0.05 mL of distilled water with the abaxial side facing upward, covered with a coverslip, and kept in the dark until imaging. Leaf epidermal tissue was examined under a macroscope to observe cellular details and PI fluorescence was excited using an HXP lamp and detected using an mRFP filter set (Excitation: BP572/25; Emission: 629/62).

### 2.5. Morphological analysis and nuclei observations of infiltrated leaves

Infiltrated leaves of *N. benthamiana* were harvested, dehydrated, embedded in Technovit® 7100 according to manufactureŕs instructions (Heraeus Kulzer, Wehrheim, Germany), sectioned to 4µm and kept at 42°C overnight to adhere the sections on glass slides. Sections were then stained either with 0.05% toluidine blue for morphology observations, or 1 µg/ml 4,6-diamidino-2-phenylindole (DAPI) (Thermo-Fischer, United States) for nuclei observations, and mounted in DPX (Sigma-Aldrich) or 90% glycerol respectively. Bright-field and epifluorescence microscopy (AxioPlan 2, Zeiss) images were respectively acquired with a digital camera (Axiocam, Zeiss, Germany).

### 2.6. Endoplasmic reticulum and the microtubule cytoskeleton observations

(A) *N. benthamiana* plants expressing HDEL-GFP (endoplasmic reticulum) and mCherry-TUa5 (cytoskeleton) were grown and agroinfiltrated as previously described. Subcellular localization of ER and tubulin was analyzed using a Zeiss LSM780 confocal microscope with 20× and 40× oil immersion objectives. HDEL-GFP was excited with a 488nm argon laser, and mCherry-TUa5 with a 561 nm DPSS laser. To control for GFP background fluorescence, non-infiltrated leaves and those infiltrated with Npp1 alone (without GFP co-expression) were used as negative controls.

### 2.7. Transmission electron microscopy analysis

Transmission electron microscopy (TEM) was performed at the Centre Commun de Microscopie Appliquée de Nice (Université Côte d’Azur) using a JEOL JEM-1400 microscope operating at 100 kV and equipped with an Olympus SIS MORADA camera. Leaf samples were fixed in 2% glutaraldehyde (in 50 mM PIPES buffer, pH 7.0, with 5 mM CaCl₂ and 0.1% tannic acid) at 4°C, then rinsed in the same buffer for 30 minutes. Post-fixation was done for 1 hour at room temperature in 1% OsO₄ and 0.8% potassium ferricyanide. Samples were rinsed in water, stained with 2% aqueous uranyl acetate for 2 hours, rinsed again, dehydrated through an ethanol series, and embedded in EPON resin. Ultrathin sections (90 nm) were collected on Formvar-coated copper grids, then contrasted with uranyl acetate and lead citrate before observation.

### 2.8. Protein immunodetection

Immunodetection of γH2A.X was performed essentially as described by De Almeida Engler et al. [60]. Plant leaves were fixed in 4% formaldehyde in 50 mM PIPES buffer (pH 6.9) at 0–4°C, dehydrated in an ethanol series, and infiltrated overnight in 50% ethanol/methacrylate mixture (BMM, butyl-methyl methacrylate 4:1). Samples were then embedded in 100% BMM with 1 mM DTT and 0.5% benzoin ethyl ether, polymerized under UV for 6 hours. Semi-thin (5 µm) sections were transferred to poly-L-lysine coated slides, dried on a hot plate and incubated overnight at 42 °C to adhere to the slides. Slides were de-plasticized in acetone (30 min), rehydrated through ethanol, and rinsed in PIPES buffer. Blocking was done in 2% BSA (30 min–2 h). Primary antibody (1:150, SAB5600038-SIGMA) was applied for 15 min at 37 °C and incubated overnight at 4 °C. Slides were then incubated 1–2 h at 37°C, washed, and incubated with anti-rabbit Alexa-Fluor 488 secondary antibody (1:300) for 2 h at 37 °C. After a final wash, slides were counterstained with DAPI (1 µg/mL), mounted in 90% glycerol, and visualized using epifluorescence microscopy (Axiocam, AxioPlan 2, Zeiss).

### 2.9. Western blotting analysis

Leaf samples were homogenized in liquid nitrogen using a Potter homogenizer. Total proteins were extracted with 2× Laemmli buffer (4% SDS, 20% glycerol, 10% 2-mercaptoethanol, 0.004% bromophenol blue, 0.125 M Tris-HCl, pH ∼6.8), supplemented with protease (cOmplete, Roche #4693132001) and phosphatase inhibitors (PhosSTOP, Roche #4906845001). Cell lysis was performed at room temperature. Lysates were centrifuged at 13,000 rpm, and supernatants were diluted 1:1 with water and boiled to denature proteins. Protein concentrations were measured using the Pierce assay (Thermo #22660) with the Ionic Detergent Compatibility Reagent (Thermo #22663). Typically, 15 µg of total protein per sample was loaded onto 15% SDS-PAGE gels (Tris-Glycine, Euromedex EU0510-B) and run at 250 V for 45 minutes. Proteins were transferred to 0.2 µm PVDF membranes (Bio-Rad) using the Trans-Blot Turbo system. Membranes were fixed in methanol and blocked in 5% BSA (in 1× TBST) for 2 hours at room temperature. Western blots were performed using the following primary antibodies: anti-phospho-RAD51 (Sigma SAB4504264, 1:500), anti-phospho-ERK1/2 (CST #9101, 1:2000), anti-RAD51 (Sigma SAB1406364, 1:500), and anti-ERK1/2 (CST #4695, 1:2000). Membranes were incubated overnight at 4 °C, followed by a 1-hour incubation with HRP-conjugated secondary antibodies (goat anti-rabbit IgG, Agrisera #AS09602; rabbit anti-mouse, Sigma A9044; 1: 10,000). PSBo (Agrisera #AS06 142-33) was used as a loading control. Detection was carried out using Immobilon Forte Western HRP substrate (Millipore WBLUF) and visualized with a FUSION FX7 imager (VILBER) using Evolution Capt Edge software. The pixel intensity ratios between the PsBO loading control and the proteins of interest were calculated using ImageJ.

### 2.10. Caspase assay

Leaf samples for caspase assays were collected at 15, 20, and 24 hours’ post-infiltration. Tissues were homogenized in liquid nitrogen and resuspended in cold 1× CASPB buffer (100 mM HEPES, 0.1% CHAPS, 1 mM DTT, pH 7.0). Lysates were centrifuged at 13,000 rpm for 10 min at 4 °C, and supernatants were either used immediately or stored at –80 °C. Protein concentrations were determined using the Pierce assay (Thermo #22660). Caspase-like activity was measured in 50 µL reactions containing 6 µg of protein and 50 µM Ac-DEVD-R110 substrate (Biotium #10226, diluted in DMSO). Fluorescence was recorded every 2.5 minutes over 4 hours at 37 °C using a LightCycler 480.

### 2.11. Reverse transcription-polymerase chain reaction (RT-PCR) analysis

Total RNA from *N. benthamiana* tissues was extracted by TRI Reagent (Sigma-Aldrich T9424), using standard procedures. RNA was resuspended in RNASE/DNASE free water. RNA concentration and purity was determined using nanodrop (Thermo scientific, NANODROP 2000) and electrophoresis. The reverse transcription kit script (Biorad, 1708891) was used for cDNA synthesize. The SYBR green PCR master mix (Takyon, MasterMix blue dTTp UF-NSMT-B0701) was used for RT-qPCR reaction. PCR were performed on 5-10ng of cDNA per well depending of the primers using CFX Opus 96 Real-time PCR instrument (#12011319) and Bio-Rad CFX Maestro software. Three reference genes were used: *Ef1a* (Elongation Factor 1), *UBQr* (Ribosomal Ubiquitin), and *GAPDH* (Glyceraldehyde 3-phosphate dehydrogenase). The candidate genes were *ATG7, BECLIN1 SOG1, ATAF1* and *WEE1* (All Primers and GENBANK accession are listed in Table S1). Considering the fact that advanced state of cell death particularly at 3 days’ post infiltration (DPI) cause general transcriptome degradation, particular attention was given to cDNA concentration deposited on the well for each samples. Between six and nine biological replicates were performed for each gene and data were subjected to statistical analyses using GraphPad Prism Software.

### 2.12. Co-immunoprecipitation experiment

For co-immunoprecipitation, *MpMIF1*, *SOG1*, and *ATAF1* coding sequences were amplified from *Nicotiana benthamiana* leaf cDNA and tagged at the C-terminus with either CMyc or HA using specific primers (All Primers and GENBANK accession are listed in Table S2). Constructs were cloned into the pH2GW7.0 vector via Gateway cloning and introduced into *Agrobacterium tumefaciens* GV3101 by electroporation. Agroinfiltration was carried out with 1:1 mixture of Prey and Bait constructs (MpMIF1-CMyc + SOG1-HA, MpMIF1-CMyc + ATAF1-HA), with empty vector used as a negative control. Infiltrated leaves were harvested at 1 and 2 DPI. Proteins were extracted from 500mg of leaf tissue homogenized in 1.5mL GTEN buffer (10% glycerol, 25 mM Tris pH 7.5, 1 mM EDTA, 150 mM NaCl, 2% PVPP), supplemented with 500 µM DTT and 0.001% Nonidet™ P-40. After centrifugation (13,000 rpm, 15 min, 4 °C), 400 µg of supernatant was incubated with 25 µL Myc-Trap® Magnetic Agarose beads (Chromotek), pre-washed in dilution buffer (10 mM Tris-HCl pH 7.5, 150 mM NaCl, 0.5 mM EDTA), for 1 hour at 4 °C. Beads were then washed three times and resuspended in 75 µL 2× SDS sample buffer, boiled for 5 min at 95 °C, and analyzed by SDS-PAGE. Typically, 40µg of total protein (Input) and 1µg of purified protein (IP) were used. Membranes were blocked with 5% milk in 1×TBST. Primary antibodies used were mouse anti-HA (Sigma H9658, 1:1000) and rat anti-Myc-TAG (Chromotek 9E1, 1:2000), incubated overnight at 4 °C. HRP-conjugated secondary antibodies (rabbit anti-mouse, Sigma A9044; goat anti-rat, Agrisera AS10 1187) were used at 1: 10,000. Signal detection followed standard chemiluminescence protocols as previously described.

### 2.13. Luciferase complementation imaging (LCI) assays

Coding sequences of *MpMIF1*, *SOG1*, and *ATAF1* were cloned from pDONR207 entry vectors into Gateway-compatible binary vectors pAMPAT-nLUC-GWY and pAMPAT-cLUC-GWY, allowing fusion to the N- and C-terminal fragments of firefly luciferase, respectively. Resulting constructs were transformed into *Agrobacterium tumefaciens* GV3101 and agroinfiltrated into *Nicotiana benthamiana* leaves as described above. At 1 and 2 days’ post-infiltration (DPI), leaves were sprayed with 1 mM D-luciferin (Sigma, L6882) in water with 0.01% (v/v) Tween-20 and incubated in the dark for 10 min. Luminescence was captured using a FUSION FX7 Imager (VILBER) equipped with a cooled camera (–80 °C), and images were processed with Evolution Capt Edge software. Six biological replicates were analyzed per interaction (two plants per replicate, two leaves per plant). Quantification of Luminescence: 4 mm leaf discs were excised and incubated in 50µL of 1 mM D-luciferin in a 96-well plate. After 10 min incubation in the dark, luminescence was measured for 1 second per well using a SAFAS Xenius XL luminometer. Each interaction was tested in 4–6 technical replicates across seven independent experiments. Data were analyzed using GraphPad Prism.

### 2.14. Statistical analysis

All experiments were independently repeated between three and nine times. Differences in expression levels, chlorophyll quantification, enzymatic activities and yH2AX ratios were tested for statistical significance by Fisher’s one-way analysis of variance by ranks (ANOVA) and the Tukey-Kramer test (Software Prism v.5.0, GraphPad).

## 3. Results

### 3.1. Quantification of cell death following co-expression of the cell death inducer Npp1 and MpMIF1

Propidium iodide (PI) staining and pictures of leaves agroinfiltrated with Npp1+MpMIF1 illustrated that MpMIF1 likely suppresses cell death at 1 and 3 DPI (Figure 1A). Control samples (Empty Vector and MpMIF1+GFP) displayed intact epidermal cells with no nuclear PI staining (Figure 1A.a,c). Similarly, Npp1+MpMIF1 co-infiltrated areas showed few PI-stained cells at 1 DPI (Figure 1A.g). In contrast, Npp1+GFP-infiltrated areas exhibited a high level of cell death, comprising 50% of nuclear PI staining as early as 1 DPI (Figure 1A.e). By 3 DPI, Npp1+GFP areas showed 100% of nuclear PI staining with apparent loss of cell wall integrity, illustrating widespread cell death (Figure 1A.f). However, in Npp1+MpMIF1 samples at 3 DPI, PI-stained nuclei were rare, and epidermal cells remained intact, resembling controls (Figure 1A.b,d,h).

**Figure 1:**
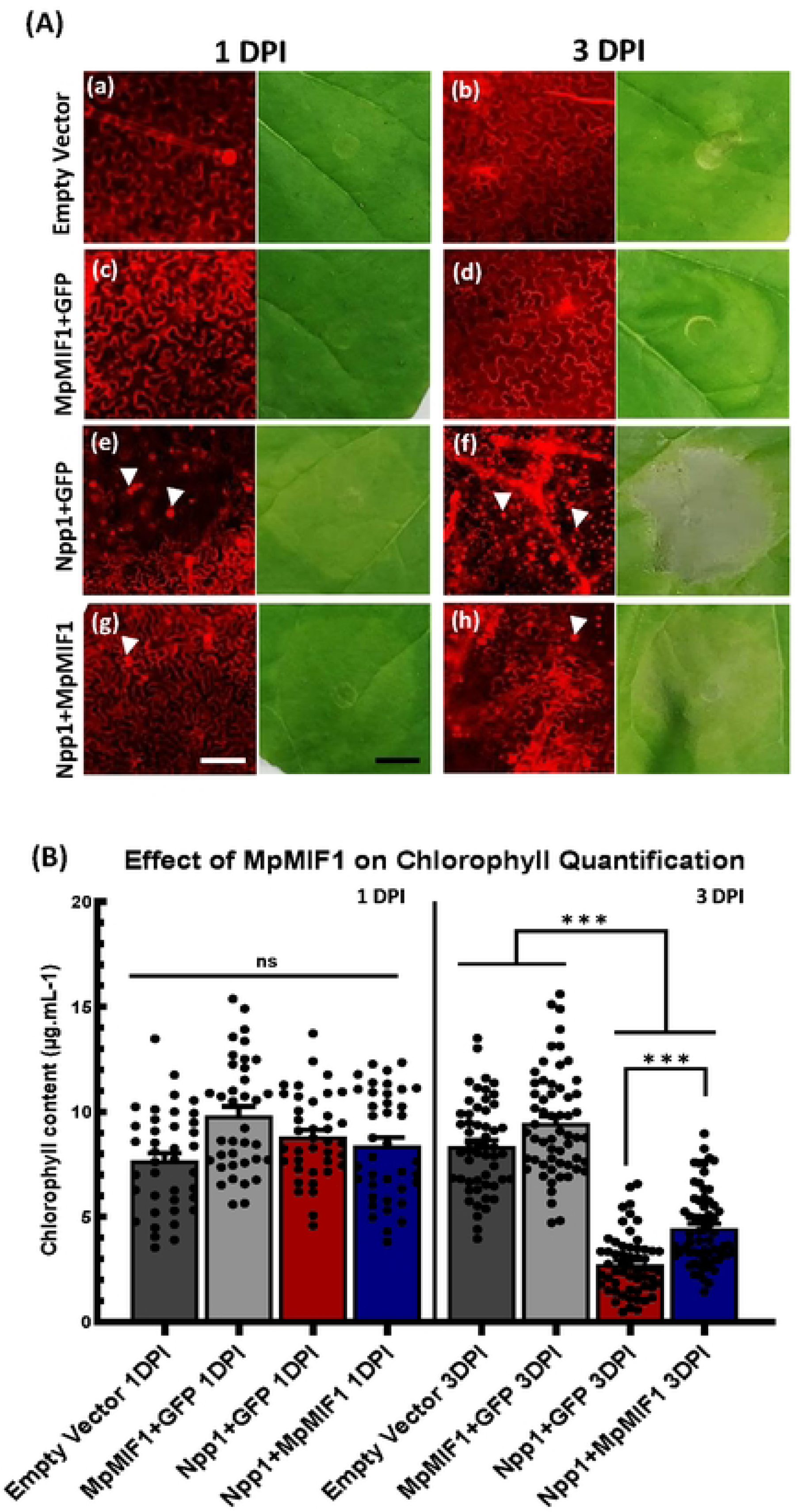
MpMIF1 prevents cell death in Nicotiana benthamiana leaves. (A) Propidium Iodide (PI) staining and leaves pictures showing cell death in N. benthamiana. Leaves were inoculated with different strains of A. tumefaciens harboring MpMIF1 construct combined with GFP construct, Npp1 combined with GFP and Npp1 combined with MpMIF1 constructs. For PI staining, the white arrowheads indicate the PI-stained nuclei of dead cells, less stained nuclei are visible in the co-infiltration of Npp1 and MpMIF1. For leaves pictures, the visible symptoms of Npp1-induced cell death are dehydration and brown lesions. These symptoms do not establish in leaf areas that express MpMIF1. Pictures were taken at 1 and 3-Days Post infiltration (DPI) with A. tumefaciens. Bars = 200 μm (B) Chlorophyll quantification in plant leaves after agroinfiltration. The total chlorophyll was measured at 1 and 3 DPI in N. benthamiana plant leaves subjected to the same treatment as above. Error bars represent the Standard Error of 12 biological replicates. The asterisks indicate statistical differences according to ANOVA and Tukey-Kramer test (***p < 0.0001, **p < 0.01, *p < 0.05, ns = not significant).

Alongside leaf infiltration observations and PI staining, chlorophyll quantification was performed in agroinfiltrated tissues since cell death is associated with chlorophyll degradation. Chlorophyll quantification showed no significant differences at 1 DPI for all treatments (Figure 1B), but by 3 DPI Npp1+GFP samples showed significantly reduced chlorophyll levels compared to Npp1+MpMIF1 (Figure 1B).

### 3.2. Morphological changes induced by Npp1-triggered cell death and MpMIF1 activity in *N. benthamiana* leaves

To assess morphological changes induced by Npp1-triggered cell death and MpMIF1 activity, we used toluidine blue and DAPI staining’s on *N. benthamiana* leaf sections (Figure 2A). In controls, leaf cells appeared intact with large central vacuoles and chloroplasts lining the plasma membrane (Figure 2A.a,b) and DAPI staining illustrated intact nuclei. Infiltration with Npp1+GFP caused visible cell damage by 1 DPI (Figure 2A.c), and by 3 DPI cells showed disrupted morphology with apparently swollen chloroplasts filling the cytoplasm, likely due to tonoplast collapse (Figure 2A.d). In contrast, co-expression with MpMIF1 led to early signs of recovery by 1 DPI (Figure 2A.e), with cells retaining their vacuole and chloroplast positioning along the cell wall. By 3 DPI, leaf tissue morphology resembled controls (Figure 2A.f). Transmission electron microscopy (TEM) analyses confirmed these observations and revealed significant morphological changes in chloroplasts among agroinfiltrated leaves (Figure 2B). Control cells from 1 to 3 DPI illustrated apparently normal organelles morphology (Figure 2B.g). In contrast, Npp1+GFP expression caused plasma membrane retraction and severe chloroplast morphological damage, illustrating distorted membranes, disorganized stroma thylakoids structures and misshaped starch grains by 3 DPI (Figure 2B.h,k,m). Notably, structures resembling enlarged plastoglobules were frequently observed in the cytoplasm in close proximity to the chloroplasts (Figure 2B.k). Co-expression with MpMIF1 diminished these effects with time illustrating altered chloroplasts at 1 DPI (Figure 2B.i) and recovered ones by 3 DPI with restored thylakoid structure, normal sized plastoglobules and restored starch grains (Figure 2B.l,m). Confocal microscopy analyses of the ER (HDEL-GFP) (Figure 3A) revealed a structured ER in control leaves, often pushed against the plasma membrane by the vacuole (Figure 3A.a, b). In contrast, the presence of Npp1 illustrated a disrupted ER architecture and cell wall integrity, leading to a less defined network along the plasma membrane in epidermal cells, with a visible marked ER disruption evident by 3 DPI (Figure 3A.c,d). Differently, MpMIF1 co-expression-maintained ER and cell wall integrity (Figure 3A.e,f). Similar confocal analyses of microtubules (MT) (mCherry-TUA5) (Figure 3B) revealed that the MT structure was clearly disturbed upon Npp1 expression causing MT bundling, visible throughout the cytoplasm and illustrating an unstructured fluorescence within stomata cells (Figure 3B. h). In the presence of MpMIF1, MT arrangement was partially restored and visible as more filamentous as in control samples (Figure 3B.g,i).

**Figure 2:**
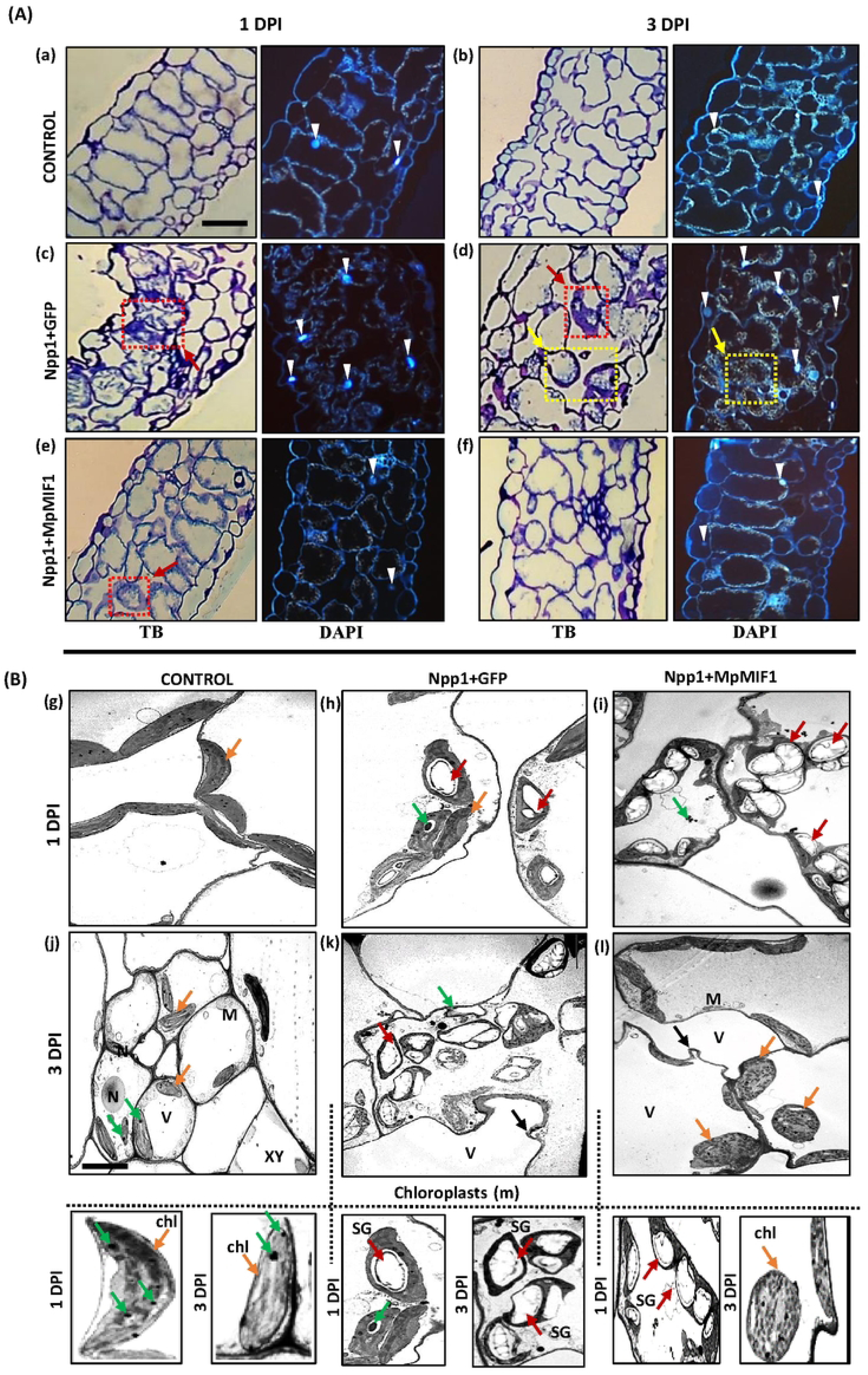
Morphological and ultrastructural analysis indicates that MpMIF1 leads to cellular recovery. (A) Toluidine blue (TB) pictures and 4’,6-Diamidino-2-Phenylindole, Dihydrochloride (DAPI) stained epidermal cells of Control (no infiltrated region), Npp1+GFP and Npp1+MpMIF1 leaf cross-sections. Observations were made at 1 and 3 DPI time points. The red arrows point to cells showing plasma membrane restrictions; the yellow arrows show enlarged chloroplasts caused by cell death induction (big white point inside plant cell) and the white triangle indicates nuclei stained by DAPI. Images were obtained using bright-field microscopy for TB and fluorescence microscopy for DAPI. Bars = 100 μm. (B) Transmission Electron Microscopy (TEM) pictures of control (non-infiltrated region), Npp1+GFP and Npp1+MpMIF1 samples at 1 and 3 DPI from N. benthamiana leaves showing subcellular structures. chl, chloroplasts (orange arrows); M, mitochondria; N, nuclei; SG, starch grains (red arrows), V, vacuole; XY, xylem cells. Black arrows point to the plasma membrane retraction; green arrows point to plastoglobules. Bars = 10 μm.

**Figure 3:**
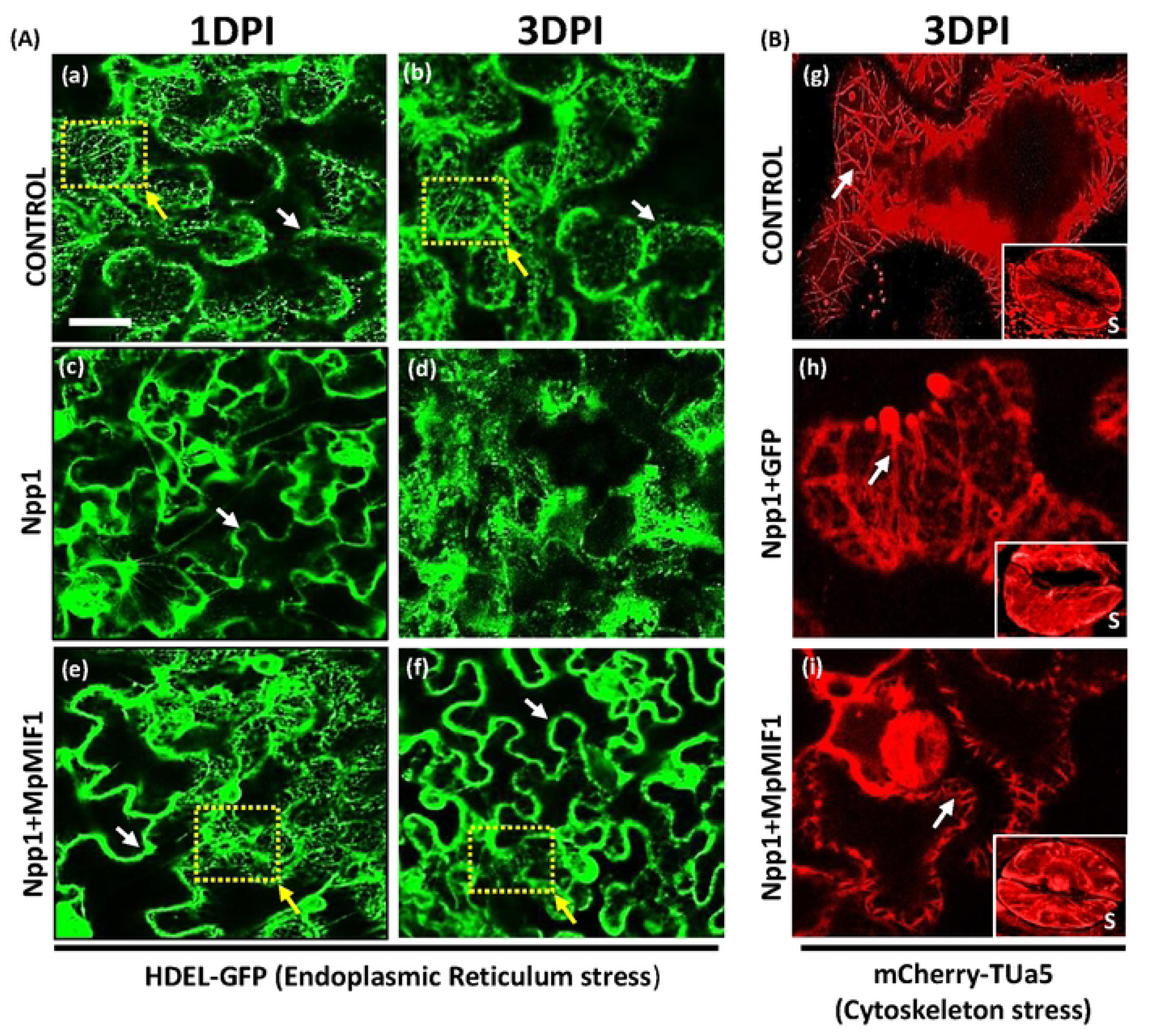
MpMIF1 protects cytoskeleton and Endoplasmic Reticulum from stress. (A) Endoplasmic reticulum structure in HDEL-GFP transformed N. benthamiana leaves. Confocal images of Control (non-infiltrated regions), Npp1+GFP and Npp1+MpMIF1 at 1 and 3 DPI. Npp1 caused disorganization of the ER in HDEL-GFP transformed N. benthamiana leaves while MpMIF1 infiltration leads to an apparent conservation of ER structure. The yellow arrows indicate cytoplasm containing ER. Bars = 100 μm. (B) Confocal pictures of mCherry-TUA5 tobacco plant showing cytoskeleton structure at 3 DPI. Npp1 damages the cytoskeleton structure of both epidermal and stomata cells leading to visible formation of larger microtubular bundles. Co-infiltration with MpMIF1 leads to a slight recovery. S letter indicates Stomata cells. White arrows indicate microtubules. Bars = 20 μm.

### 3.3. MpMIF1 appears to be involved in several plant cell death pathways

To investigate the potential involvement of MpMIF1 in known plant cell death pathways, we selected markers of autophagy, senescence and apoptosis-like activities as represented in Figure 4A. In addition, we investigated DNA DSBs, as they are closely associated with various active cell death processes among all organisms [61], as well as DNA repair and a cell cycle checkpoint control marker (Figure 4A).

**Figure 4:**
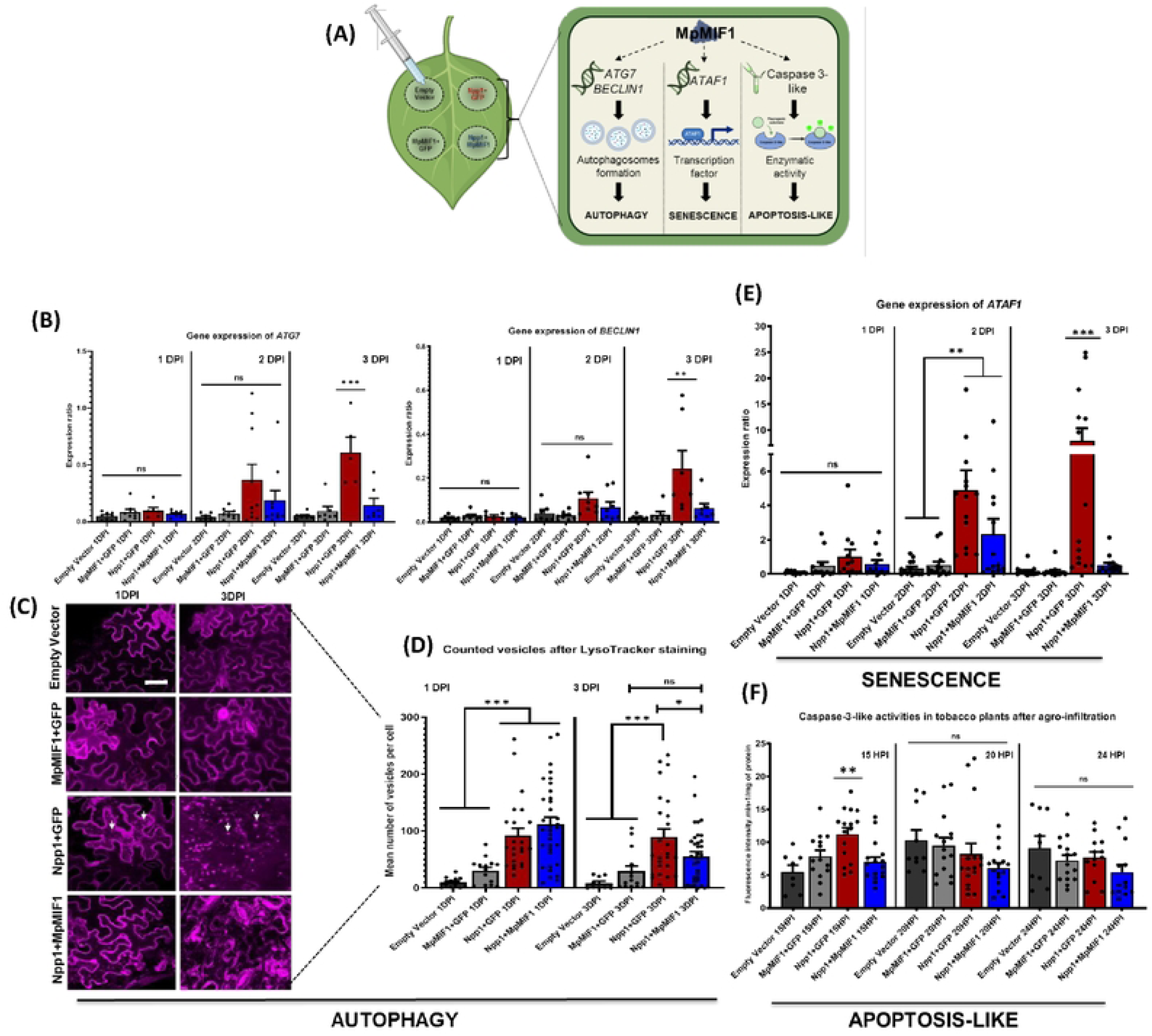
MpMIF1 impairs plant cell death genes and functions. (A) Schematic representation of the strategy used to study the impact of MpMIF1 on selected plant cell death candidates. (B) Expression ratio of autophagy-related transcripts ATG7 and BECLIN1 in agroinfiltrated leaf areas at 1, 2 and 3 DPI. (C) LysoTrackerTM Red DND-99 staining on N. benthamiana leaves after agroinfiltration. Representative pictures of confocal microscopy at 1 and 3 DPI for all the treatments. The white arrows point to fluorescent vesicles assimilated to autophagosomes. (D) Diagram showing the number of fluorescent vesicles per cell for the different treatments at 1 and 3 DPI. MpMIF1 decreases the number of autophagosomes per cell. (E) Expression ratios of senescence-related-gene ATAF1 at 1, 2 and 3 DPI. (F) Caspase 3-like activity at 15, 20 and 24 hours’ post-infiltration (HPI). For RT-QPCR expression analyses, expression ratios are shown relative to the expression of the house-keeping genes GAPDH, EF1a and Ribosomal Ubiquitin. For all diagrams, error bars represent the Standard Error of 6 to 9 biological replicates. The asterisks indicate statistical differences according to ANOVA and Tukey-Kramer test (***p < 0.0001, **p < 0.01, *p < 0.05, ns = not significant).

#### 3.3.1. Autophagy pathway

The effect of MpMIF1 on autophagy was investigated by analyzing the expression of the autophagy-related genes *ATG7* and *BECLIN1*, which are essential for autophagosome formation [37,38]. At 3 DPI, both genes were significantly upregulated in Npp1+GFP samples, while expression remained low in Npp1+MpMIF1 and controls (Figure 4B). Using LysoTracker staining, specific for acidic organelles such as autophagosomes [62], we observed increased vesicle formation at 1 DPI in both Npp1+GFP and Npp1+MpMIF1 (Figure 4C). By 3 DPI, vesicle number remained high in Npp1+GFP but decreased in Npp1+MpMIF1, suggesting that MpMIF1 modulates autophagy activity (Figure 4D).

#### 3.3.2. Senescence pathway

We examined the effect of MpMIF1 on senescence-induced cell death by measuring *ATAF1* gene expression using RT-qPCR. *ATAF1* is a known positive regulator of senescence in plants [54]. At 1 DPI, *ATAF1* levels increased at 2 DPI in both Npp1+GFP and Npp1+MpMIF1. By 3 DPI, *ATAF1* transcript levels remained high in Npp1+GFP but returned to control levels in Npp1+MpMIF1, indicating MpMIF1 may counteract prolonged senescence signaling (Figure 4E).

#### 3.3.3. Apoptosis-like pathway

To assess caspase-3-like activity, we used the Ac-DEVD-AMC substrate, which is specific for caspase-3 in mammals [63]. Since caspase-3 is typically activated during early apoptosis, we analyzed its activity at 15, 20, and 24 hours post-infiltration (HPI). At 15 HPI, Npp1+GFP samples showed elevated caspase-3-like activity (11.2 fluorescence intensity·min⁻¹·mg⁻¹), while Npp1+MpMIF1 samples showed reduced activity (6.9 fluorescence intensity·min⁻¹·mg⁻¹), similar to controls. No differences were observed at later time points (Figure 4F). This suggests that MpMIF1 suppresses early apoptosis-like activity. To confirm the specificity of this fluorescent signal, we attempted to use the caspase-3 inhibitor Ac-DEVD-CHO. Despite multiple attempts, no significant differences were detected with or without the inhibitor (Figure S2). This result is not entirely unexpected, given that caspase-like activity in plants can arise from various proteases, such as caspases or vacuolar processing enzymes (VPEs), which may interact differently with the substrate [50,63].

### 3.4. MpMIF1 is involved in several DNA damage response mechanisms

#### 3.4.1. DNA double strand breaks (DSBs)

The potential DSBs induced by Npp1 and the potential involvement of MpMIF1 was examined via immunolocalization of the phosphorylated form of H2AX histone, a marker of DSBs (γH2A.X) (Figure 5A). We analyzed 50 sections per sample to calculate the ratio of γH2A.X-positive nuclei (fluorescent green) and observed nuclei by DAPI-staining (Figure 5C). At 1 DPI, 75% of nuclei were γH2A.X-positive in Npp1+GFP samples, while only 38% were positive in Npp1+MpMIF1 samples and similar to the control (Figure 5C), suggesting that MpMIF1 reduces DNA damage under cell death conditions induced by Npp1. No significant differences were seen at 2 DPI, due to advanced cell death state of Npp1+GFP sample.

**Figure 5:**
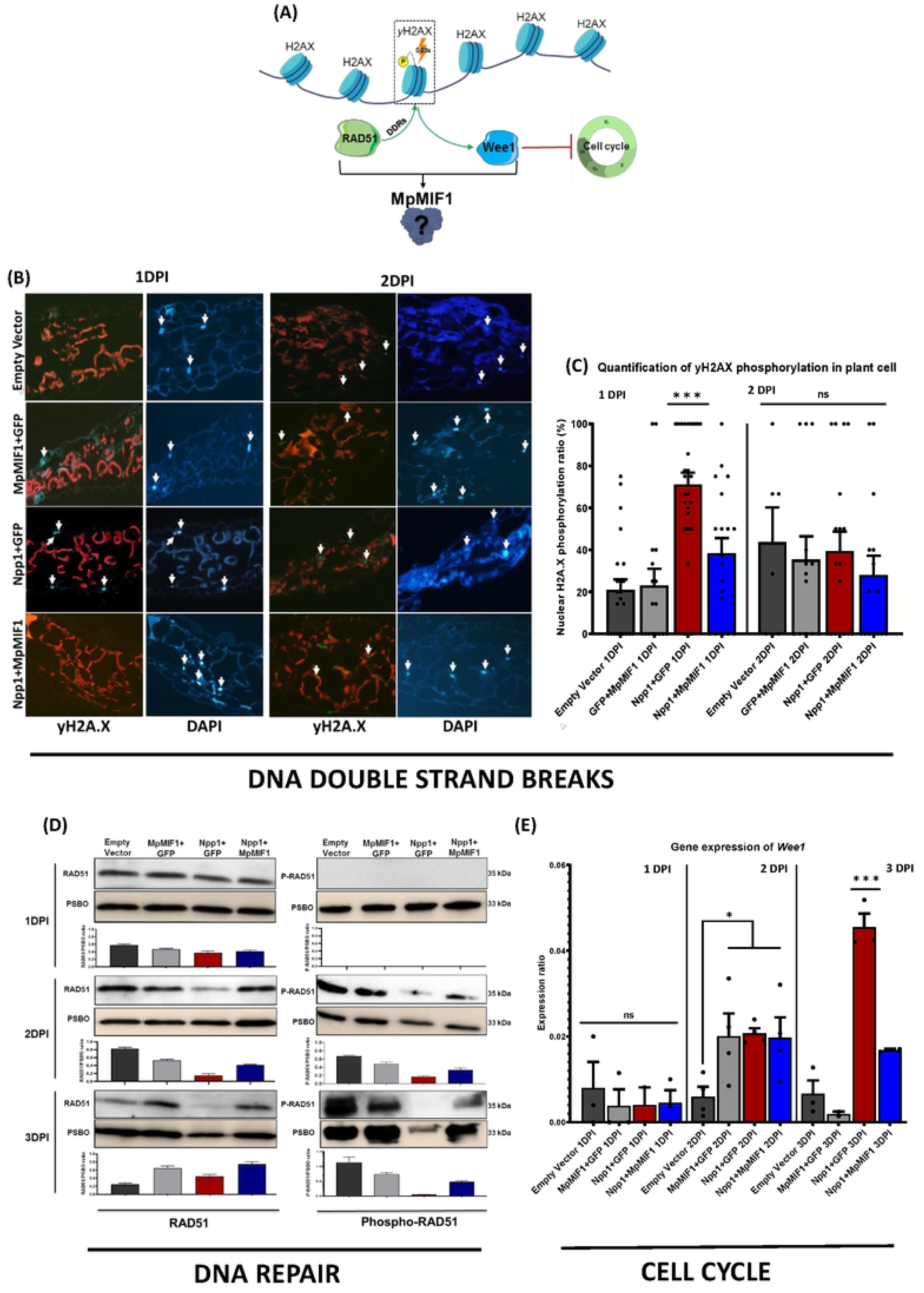
MpMIF1 impairs important DNA Damage Response genes conserved between plants and mammals. (A) Schematic representation of DNA Damage Response (DDR) mechanisms studied with MpMIF1. When DDR occurs, yH2AX attracts DNA repair proteins such as RAD51. In parallel, in order to avoid DNA replication mistakes, WEE1 is activated to stop the cell cycle in G2/M phase for genomic integrity checking before entry into mitosis. (B) Immunolocalization of yH2AX (green nuclei) in N. benthamiana agroinfiltrated leaves at 1 and 2 DPI. Red fluorescent cell walls result from auto-fluorescent signal of the Dual Band pass filters. The white arrows indicate nuclei inside the plant cells. Green fluorescent nuclei are yH2AX positive and contain DNA damages. Bars = 100 μm. (C) Quantification of H2A.X phosphorylation in plant cells at 1 and 2 DPI, shown as a percent of Green fluorescent nuclei (yH2AX positive) over DAPI nuclei (total nuclei number). (D) Western Blot analysis of RAD51 and Phospho-RAD51 at 1,2 and 3 DPI for all treatment. The band at 35kDA represents RAD51 (Upper blot), the band at 33Kda (lower blot) represent PSBo Oxygen-evolving enhancer protein 1 protein used as protein loading control to compare the total protein quantity. Graphical analysis below blots for each sample represent the ratio between either RAD51 (left) or P-RAD51 (right) on PSBO loading control. Error bars represent the Standard Deviation. MpMIF1 maintains RAD51 phosphorylation and protein quantity during cell death stress response. (E) Expression ratio of WEE1 in agro-infiltrated leaf areas at 1, 2 and 3 DPI. WEE1 overexpression is correlated to cell cycle arrest. Asterisks indicate statistical differences according to ANOVA and Tukey-Kramer test (***p < 0.0001, **p < 0.01, *p < 0.05, ns = not significant).

#### 3.4.2. DNA repair

To assess the impact of MpMIF1 on DDR, we analyzed the presence and abundance of RAD51 known to be implicated in DNA repair and its phosphorylated active form (Figure 5A) [64] using immunoblotting. At 1 DPI, RAD51 levels were similar amongst treatments. At 2 and 3 DPI, RAD51 ratio levels decreased in Npp1+GFP but remained stable in Npp1+MpMIF1 (Figure 5D). Detection of phosphorylated RAD51 revealed a clear positive signal in NPP1+MpMIF1 samples at 2 and 3 DPI, whereas only a weak or no signal was observed in NPP1+GFP samples (Figure 5D). Control samples also showed phospho-RAD51 detection, indicating that agroinfiltration can trigger DDR. To further investigate MpMIF1’s role in DDR, we measured *RAD51* gene expression. However, no significant differences were detected (data not shown). These results suggest that MpMIF1 helps to preserve RAD51 protein levels and activity, contributing to DNA repair without upregulating *RAD51* expression.

#### 3.4.3. Cell cycle checkpoint

Upon DNA damage, both plant and animal cells activate checkpoint control pathways, which delay cell cycle progression to allow DNA repair enzymes to correct anomalies. One key gene involved in DNA damage checkpoint control is *WEE1*, which encodes a protein kinase that regulates the G2-to-M transition by inhibiting cell cycle progression (Figure 5A) [65]. Gene expression analysis of *WEE1* revealed a significant overexpression in the Npp1+GFP treatment at 3 DPI, but not in Npp1+MpMIF1 samples (Figure 5E). No significant differences were detected at 1 DPI, while a general increase in *WEE1* expression was observed at 2 DPI among all co-expressed samples, likely due to the effect of agroinfiltration (Figure 5E). These results suggest that MpMIF1 contributes to maintain normal cell cycle control following DNA damage, even in the presence of a cell death inducer.

### 3.5. Conservation of MIF signaling pathways between mammals and plants

In addition to investigating the effect of MpMIF1 on known plant cell-death markers, we investigated two of the major pathways affected by MIF in mammals that are conserved in plants. We focused on the MAPK and the SOG1 pathways (Figure 6A).

**Figure 6:**
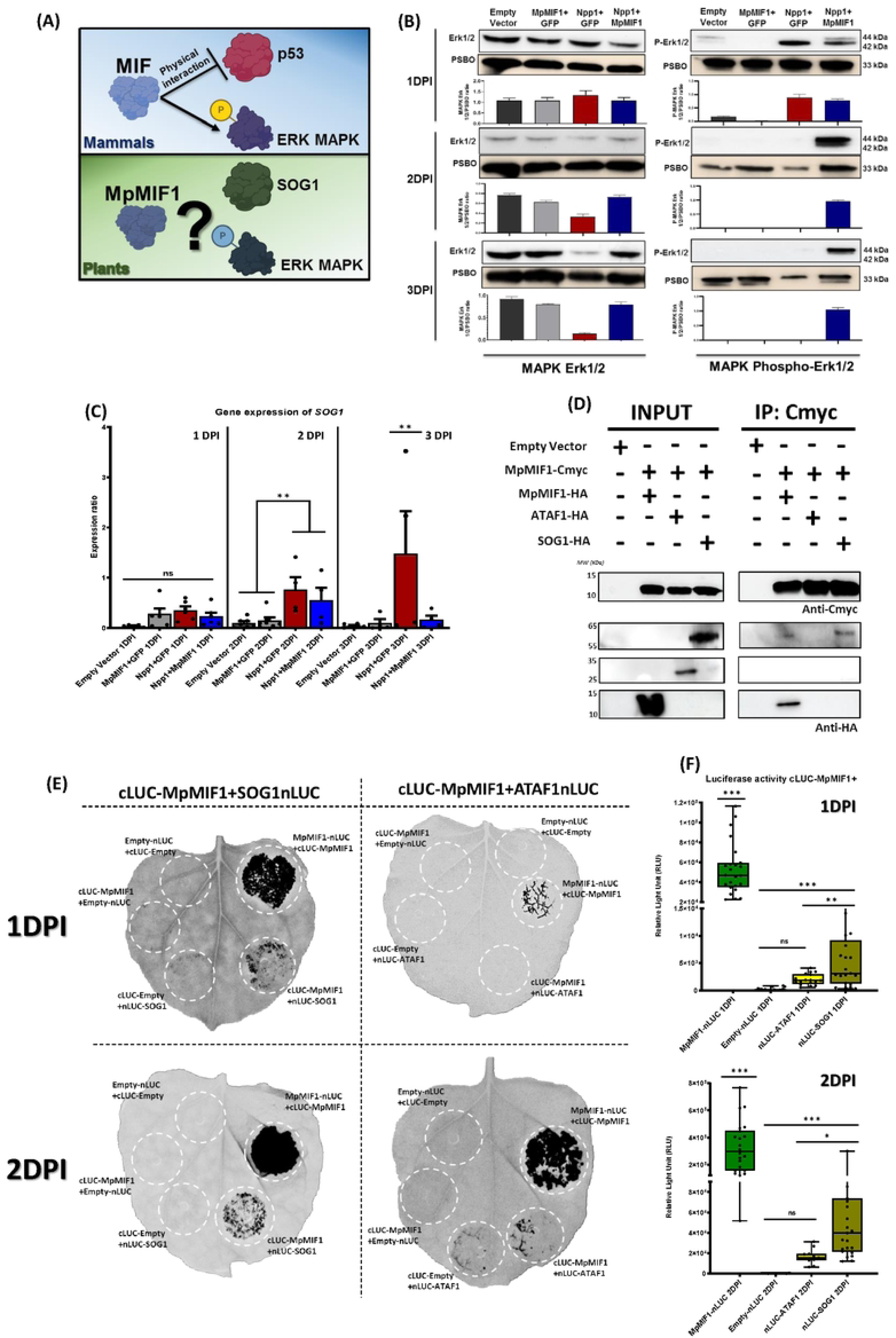
Conservation of MIF signaling pathways between mammals and plants. (A) Schematic representation of two major MIF targets in mammals and their potential counterparts in plants. (B) Western Blot analysis of MAPK Erk 1/2 and MAPK Phospho-Erk1/2 at 1,2 and 3 DPI for all treatments. The 42/44kDa band corresponds to MAPK Erk1/2 (Upper blot) and the 33Kda band (lower blot) corresponds to PSBo Oxygen-evolving enhancer protein 1 protein used as a loading control. Erk1/2 MAPK are strongly phosphorylated in the presence of MpMIF1. Graphical analysis below blots for each samples represent the ratio between either MAPK Erk1/2 (left) or MAPK Phospho-Erk1/2 (right) on PSBO loading control. Error bars represent the Standard Deviation. (C) Expression ratios of SOG1 in agroinfiltrated leaf areas at 1, 2 and 3 DPI. Total mRNA from leaves exposed to specific treatments were used for assessing transcript abundance relative to transcripts of control genes GAPDH, EF1a and Ribosomal Ubiquitin. Npp1 induces strong overexpression of SOG1 while MpMIF1 counteracts this overexpression. (D) Co-immunoprecipitation of epitope-tagged SOG1 and ATAF1. Tandem of MpMIF1-CMyc/SOG1-HA and MpMIF1-CMyc/ATAF1-HA were transiently co-expressed in N. benthamiana. At 1-2 dpi, proteins were extracted and immunoprecipitation was performed for the CMyc tag proteins using Myc-Trap® Magnetic Agarose. MpMIF1-CMyc protein was detected with an α-CMyc antibody and HA tag proteins with a monoclonal α-HA antibody. Ponceau staining was used as a loading control for the input. Co-immunoprecipitation was repeated three time with similar results. (E) Split-luciferase assay showing in planta interaction between MpMIF1 and SOG1 proteins. ATAF1 protein was also studied as a NAC domain protein involved in senescence and transcriptionally affected by MpMIF1. cLUC-MpMIF1 was transiently co-expressed at 1 and 2 DPI with SOG1-nLUC, ATAF1-nLUC or EV-nLUC (empty vector) in N. benthamiana. Reverse nLUC and cLUC construction between Prey and Bait was also tested (Figure S4). Black spots correspond to detected luminescence signal under CCD-camera to show luciferase activities. (F) Box plot showing luciferase activity from the split-luciferase complementation assay using Luminometer. The luminescence from seven independent experiments was quantified in single 4 mm punch disks placed into 96 well plate and incubated with 1mM luciferin. For all diagrams, error bars represent the Standard Error and the asterisks indicate statistical differences according to ANOVA and Tukey-Kramer test (***p < 0.0001, **p < 0.01, *p < 0.05, ns = not significant).

#### 3.5.1. MpMIF1 affects the plant MAPK pathway

Western blotting was performed to examine Erk1/2-like phosphorylation in the presence of MpMIF1 following cell death induction by Npp1. To confirm the presence of total Erk1/2 protein, blots detecting the normal (non-phosphorylated) form of Erk1/2 (Erk1/2) were analyzed (Figure 6B). Western blot ratio analysis showed that total Erk1/2 protein levels declined in Npp1+GFP samples over time, suggesting degradation due to cell death (Figure 6B). In contrast, the phosphorylated form (p-Erk1/2) remained detectable at 2 and 3 DPI only in Npp1+MpMIF1 samples, indicating that MpMIF1 sustains MAPK activation under cell death conditions induced by Npp1 (Figure 6B).

#### 3.5.2. MpMIF1 affects SOG1 expression

At 1 DPI, no significant differences in *SOG1* expression were observed among the four treatments (Empty vector, MpMIF1 + GFP, Npp1 + GFP, and Npp1 + MpMIF1) (Figure 6C). By 2 DPI, *SOG1* expression increased in both Npp1+GFP and Npp1+MpMIF1 samples (Figure 6C). At 3 DPI, *SOG1* remained highly overexpressed in Npp1+GFP samples, whereas in Npp1+MpMIF1 samples, its expression returned to control levels (Figure 6C). These results suggest that MpMIF1 prevents (directly or indirectly) the strong upregulation of *SOG1* in general associated with cell death.

#### 3.5.3. MpMIF1 appears to bind to SOG1

In mammals, MIF physically interacts with p53 to suppress its pro-apoptotic activity [22–24]. Given that SOG1 is considered a functional analog of p53 in plants, we investigated the potential physical interaction between MpMIF1 and SOG1. Additionally, we examined the potential binding to ATAF1, a member of the NAC domain protein family (NAM, ATAF1/2, and CUC2), to determine whether MpMIF1 could specifically interact with SOG1 or could target multiple NAC proteins. For that, co-immunoprecipitation (Co-IP) and split luciferase complementation assays were performed. We transiently co-expressed epitope-tagged (*CMyc* and *HA*) versions of *MpMIF1*, *SOG1*, and *ATAF1* in *N. benthamiana* and performed Co-IP assays using anti-CMyc magnetic beads. Prior to agroinfiltration, we confirmed that MpMIF1-TAG retained its function of cell death inhibition (Figure S3). As MpMIF1 forms homotrimers, co-expression of MpMIF1-CMyc and MpMIF1-HA served as a positive control (Figure 6D). Immunoblotting with monoclonal CMyc and HA antibodies detected a SOG1-HA pull-down between 55-65 kDa with MpMIF1-CMyc, whereas no interaction was observed for ATAF1-HA (Figure 6D). A faint band was also present at the same molecular weight for MpMIF1-CMyc and MpMIF1-HA, which represents the dimeric form of MpMIF1-HA associated with a MpMIF1-CMyc monomer (Figure 6D).

As a second approach, we used a split luciferase assay, in which proteins of interest were translationally fused to the N-terminal (nLUC) and C-terminal (cLUC) fragments of firefly luciferase. Successful interaction resulted in luciferase fragments complementation, which can be quantified by luminescence. As a positive control, we co-expressed MpMIF1-nLUC with cLUC-MpMIF1. As a negative control, we co-expressed tagged MpMIF1, SOG1, and ATAF1 with empty vector controls. Co-expression of cLUC-MpMIF1 with SOG1-nLUC resulted in strong luciferase activity at 1 and 2 DPI (Figure 6E), confirmed by both imaging and luminescence quantification (Figure 6F). Similarly, co-expression of MpMIF1-nLUC with cLUC-SOG1 yielded significant luciferase activity at 1 DPI compared to the negative control (cLUC-Empty) (Figure S4). However, at 2 DPI, this luminescence was not significantly different from the negative control (Figure S4). This result can be explained by differences in 5’ and 3’position of cLUC and nLUC in the pAMPAT-nLUC-GWY and pAMPAT-cLUC-GWY plasmids. In contrast, co-expression of cLUC-MpMIF1 or MpMIF1-nLUC with ATAF1-nLUC or cLUC-ATAF1 resulted in weak or undetectable luciferase activity (Figure 6E). Our results from both Co-IP and split luciferase assays in *N. benthamiana* strongly suggest a direct physical interaction between MpMIF1 and SOG1, but not MpMIF1 and ATAF1.

## 4. Discussion

Our results demonstrate that MpMIF1 contributes to the regulation of plant cell death and immunity by inhibiting key factors involved in the activation of various cell death pathways and by maintaining DNA repair mechanisms. In 1999, Hudson et al. [22] showed that human MIF interacts with p53 and inhibits its activity [22]. Based on this observation, we examined the potential interaction between MpMIF1 and SOG1, the functional equivalent of p53 in plants and a master regulator of DDR and cell death [51,52]. Interestingly, this study provides evidence that MpMIF1, is capable of interacting with central regulators of cell death and DDR in plants, like the MIF-p53 interaction observed in animals [22–24]. This information is particularly relevant given that aphid infection is known to cause substantial cellular damage in host plants [6,66]. Our supplementary data (Figure S5) further confirmed that *in vivo* aphid infection followed by sudden parasite withdrawal affects multiple markers of cellular stress associated with cell death, most notably the expression of SOG1. These results are consistent with and strongly support the role of SOG1 in regulating plant PCD and DDR during aphid infection.

### 4.1. MpMIF1 plays a protective role preventing cell death, safeguarding organelles from stress-related damage potentially promoting cellular recovery mechanisms

Morphological analysis of agroinfiltrated leaves and PI staining revealed that tissue damage was reduced when MpMIF1 was co-expressed with the cell death inducer Npp1, confirming both the ability of Npp1 to trigger cell death in *Nicotiana benthamiana* [57,58], and the capacity of MpMIF1 to counteract this effect. As well, chlorophyll content in Npp1+MpMIF1 samples was significantly reduced compared to the Empty Vector and MpMIF1+GFP controls, confirming that MpMIF1 can only partially inhibit cell death [8].

Other hallmarks were vacuolar cell death when Npp1 was expressed, including plasma membrane retraction and apparent tonoplast rupture. Tissue analysis by TEM provided a more detailed characterization of cell death features and further revealed the modulatory effect of MpMIF1. Temporal observations, such as at 3 DPI illustrated strong signs of cellular recovery when co-expressing MpMIF1 on leaf cells with the cell death inducer Npp1. A significant reduction in starch grains size was observed at 3 DPI, suggesting mobilization of energy reserves to support recovery. Indeed, studies highlighted that the number, size, and ultrastructure of chloroplasts, including thylakoids, starch grains, and plastoglobules serve as reliable indicators of plant stress responses [67]. Our observations have illustrated normal sized plastoglobules in Npp1+MpMIF1 at 3 DPI versus visible increased sized in Npp1+GFP samples. Changes in plastoglobules number and size are associated with both abiotic (e.g., drought, light stress) and biotic (e.g., pathogen infection) stresses [67,68]. Beyond their role in photosynthesis, chloroplasts are central for plant defense and PCD, acting as a key source of signaling molecules such as reactive oxygen species (ROS) and contributing to the biosynthesis of salicylic acid and jasmonic acid [35].

ER and MTs structures in *N. benthamiana* leaves were monitored upon stress caused by Npp1 and potential recovery induced by MpMIF1. Our results demonstrated that co-expression of Npp1 and MpMIF1 maintained a more structured ER similar to uninfiltrated cells. ER was present around the plasma membrane and nucleus, indicating preserved integrity, compared to the disrupted ER observed in Npp1+GFP samples at 3DPI. ER stress is triggered by the accumulation of misfolded proteins under adverse conditions activating unfolded protein response (UPR) eventually leading to apoptosis in mammals [69].

Interestingly, previous studies suggested a link between ER stress, caspase-like activity, autophagy, and vacuolar cell death [69,70]. For example, infection of *Arabidopsis* roots by *Piriformospora indica* causes tonoplast rupture, cytoplasmic lysis, and ER swelling highlighting an important role of the ER in plant innate immunity [69,70].

Regarding the cytoskeleton, co-expression of Npp1+GFP led to notable cytoskeletal damage, including the formation of MT bundles in both palisade parenchyma and stomatal leaf cells (Figure 3H). However, co-infiltration with MpMIF1 led to partial recovery of MT architecture revealing less thick bundled MTs (Figure 3I). It is well established that cytoskeletal remodeling, affecting both microtubules and actin filaments, is a hallmark of PCD in plants [71]. Thus, our cellular structure observations strongly suggest that MpMIF1 contributes to the recovery of chloroplasts, as well as ER and MT architecture, under stress conditions such as ectopic cell death induction.

### 4.2. MpMIF1 modulates genes involved in cell death and DNA damage response in plants

As sessile organisms, plants are continuously exposed to environmental stresses that can cause DNA damage [30]. Unable to avoid unfavorable conditions, plants have evolved sophisticated and tightly regulated mechanisms to control PCD and DNA repair processes [72]. To assess whether MpMIF1 could affect these pathways, we examined several marker genes associated with PCD and DNA repair pathways.

The expression of autophagy-related genes ATG7 and BECLIN1 showed a high overexpression at 3 DPI in Npp1+GFP leaf samples, while their expression remained close to the control in the Npp1+MpMIF1 condition (Figure 4B). Since ATG7 and BECLIN1 are key regulators of autophagosome formation and maturation [37,38], their expression patterns align with the increased number of vesicles labeled by LysoTracker Red DND-99 observed in the Npp1+GFP condition at 3 DPI. These findings support the idea that MpMIF1 inhibits excessive autophagy induction. In the absence of ATG genes, the HR-related PCD has been shown to spread beyond infection sites to neighboring and even distant tissues, underscoring the role of autophagy in controlling plant immune responses [73,74]. Additionally, ER stress was shown here to be avoided by MpMIF1 and has been previously reported to trigger autophagosomes formation in *Arabidopsis* seedlings [69]. Those data are consistent with the observed decrease in the number of autophagosomes in agroinfiltrated Npp1+MpMIF1 samples at 3 DPI. Moreover, recent studies have demonstrated that MIF can regulate autophagy in mammals as well [75]. Altogether, this suggests that aphid MpMIF1 may suppress autophagy to help maintaining cellular homeostasis during infection.

In plants, cellular senescence is a tightly regulated process that recycles nutrients and aids stress adaptation [76]. To assess the role of MpMIF1 in senescence-associated cell death, we measured the expression of *ATAF1*, a positive senescence regulator [54]. We showed here that MpMIF1 modulates senescence signaling maintaining *ATAF1* at a low level at 3 DPI, unlike the high expression seen in controls. Mammalian MIF similarly inhibits senescence-related genes [77], supporting a comparable role for MpMIF1 in plants. Since chloroplast degradation and reduced photosynthesis are early senescence signs, [78], our TEM data illustrating preserved thylakoids at the presence of MpMIF1 suggest that it likely prevents senescence. Furthermore, autophagy-related genes (*ATGs*) have been shown to be involved in senescence [78], and our data suggest that MpMIF1 may also affect senescence by repressing autophagy signaling.

Caspase-like activities play a central role in most PCD forms [79]. Although classical caspase genes are absent in plants, DEVDase activity, a marker for caspase-3-like function, has been reported during various plant cell death processes [63]. In our study, we found a significant increase in caspase-like activity upon Npp1+GFP agroinfiltration at 15 HPI, while its activity in Npp1+MpMIF1 remained similar to the untreated control (Figure 4F). This suggests that MpMIF1 suppresses early activation of caspase-3-like enzymes during cell death onset. To assess whether Npp1 induces DSBs and the potential protective role of MpMIF1, we performed immunolocalization of the phosphorylated histone variant γH2A.X, a conserved DSBs marker [46,72]. At 1 DPI, Npp1+GFP samples showed numerous γH2A.X-positive nuclei (green fluorescence), while Npp1+MpMIF1 samples illustrated reduced labelled nuclei, suggesting that MpMIF1 prevents DSBs in presence of a cell death inducer.

In animals, apoptosis is associated with the accumulation of γH2A.X [72]. Phosphorylation of H2A.X is a key step in initiating the DNA DDR, as γH2A.X recruits repair factors like RAD51 and BRCA1 to sites of DSBs [72]. DDR restores genomic stability after DSBs, with RAD51 playing a crucial role in homologous recombination repair by promoting homologous pairing and strand exchange [47]. Our western blot data illustrating RAD51 phosphorylation suggests that MpMIF1 supports DDR during cell death conditions induced by Npp1. Interestingly, mammalian studies describe a MIF–BRCA1–RAD51 axis involved in homologous recombination and ferroptosis cell death [80].

In response to DNA damage, both plants and animals activate checkpoint control pathways that delay cell cycle progression to allow repair [50]. WEE1, a kinase that inhibits cell cycle progression by phosphorylating and inactivating CDKs is key in this process [55,56]. Our RT-qPCR data showed a strong *WEE1* upregulation at 3 DPI in Npp1+GFP-treated leaf cells, indicating induced cell cycle arrest. This upregulation was absent in Npp1+MpMIF1 samples, suggesting that MpMIF1 helps maintaining cell cycle progression under reduced stress conditions. As well, *WEE1* is activated by ATR and ATM kinase signaling in response to DNA damage, so its overexpression in our cell death Npp1+GFP-treated leaf cells indicates DNA damage checkpoint activation [56,65]. Thus, our findings suggest that MpMIF1 modulates DNA damage sensing (γH2A.X), repair (RAD51), and cell cycle checkpoint (*WEE1*) control under genotoxic stress.

### 4.3. Conservation of MIF signaling pathways across kingdoms

In addition to studying MpMIF1 effects on known plant cell death markers, we investigated conserved cell death-related pathways regulated by MIF in mammals. Specifically, we focused on the MAPK pathway and the transcription factor SOG1, considered a functional analog of mammalian p53 in plants [51,52]. Mammalian MIF is highly pleiotropic regulating the expression of 65 genes and participating in 24 protein–protein interactions [81]. However, one of its key functions is to promote ERK1/2 (p44/p42 MAPKs) phosphorylation via the CD74 receptor [25], which enhances cell proliferation and inhibits PCD. The MAPK cascade, conserved in plants, regulates both defense responses and cell proliferation, including the HR and PCD [82,83]. Our data show that MpMIF1 enhances and sustains ERK1/2-like phosphorylation under cell death conditions induced by Npp1. In animals, prolonged ERK signaling not only supports apoptosis but also cell cycle progression, regeneration, and differentiation [84]. Since MpMIF1 restricts PCD in plants, its activation by ERK1/2 may help to promote cell survival and maintain homeostasis in response to cell conditions induced by Npp1. In mammals, ERK1/2 activation suppresses caspase activity [85], and increase RAD51 protein level and stability [86]. Similarly, we showed that MpMIF1 modulates caspase-like activity and RAD51 stability in plants. As well, γH2AX is also essential for sustained ERK1/2 activation, which can lead to p53 phosphorylation following DNA damage [67,68,87]. This suggests that ERK1/2 is a central hub linking DDR and PCD across kingdoms. Since plants lack MIF receptors such CD74, CXCR2, or CXCR4 [88], our results indicate that MpMIF1 triggers ERK1/2 via a distinct, plant-specific mechanism involving interactors to be identified.

In mammals, p53 regulates PCD by controlling genes involved in cell cycle arrest, apoptosis, senescence, and DNA repair [89]. MIF inhibits both the expression and activity of p53, thereby reducing apoptosis [22–24]. This interaction is also seen in invertebrates, where MIF from *Biomphalaria glabrata* promotes immune cell proliferation and suppresses p53-dependent apoptosis [90]. While plants lack a direct p53 homolog, the transcription factor SOG1 is considered its functional equivalent [51,52]. We examined the effect of MpMIF1 on *SOG1* expression and found that *SOG1* was upregulated at 2 DPI in both Npp1+GFP and Npp1+MpMIF1 samples. By 3 DPI, *SOG1* remained high in Npp1+GFP samples but returned to baseline in Npp1+MpMIF1. These findings suggest that MpMIF1 modulates SOG1 by maintaining moderated expression levels to support DNA repair, whereas high SOG1 expression, in presence of Npp1 may promote cell cycle arrest and cell death. In mammals, p53 activity varies depending on the cellular response, being more strongly regulated during PCD than during DDR [89]. This is consistent with the higher SOG1 expression observed in Npp1+GFP condition compared to Npp1+MpMIF1 at both 2 and 3 DPI. Notably, SOG1 has also been shown to regulate WEE1 transcription, while ATR-WEE1 signaling promotes SOG1 translation, forming a positive feedback loop essential for fine-tuning the DDR [56]. These data are consistent with the overexpression of SOG1 and WEE1 observed in Npp1+GFP samples at 3 DPI. Although SOG1 is primarily activated via ATM/ATR signaling, alternative pathways, such as MAPK cascades have also been proposed to regulate its activity [65]. Our findings on ERK1/2 phosphorylation by MpMIF1 further support a link between MAPK signaling and SOG1 regulation. To further investigate the MpMIF1–SOG1 regulatory mechanisms, co-immunoprecipitation and split-luciferase complementation assays confirmed that MpMIF1 physically interacts with SOG1 in planta. MpMIF1-CMyc co-precipitated with SOG1-HA, and co-expression of cLUC–MpMIF1 with SOG1–nLUC produced a strong luminescence. This interaction may explain how MpMIF1 modulates SOG1-driven PCD, resembling the MIF–p53 regulatory pathway in mammals.

## 5. Concluding Remarks

Herein, we demonstrate that MIF1 from *Myzus persicae* interacts with SOG1, the central regulator of DDR pathway in plants, and impacts all examined cell death and DDR pathways. Based on our findings, we propose a model (Figure 7) integrating candidate genes involved in the molecular network governing cell death and DDR, likely triggered by tissue damage from aphid stylet penetration and the subsequent release of MpMIF1 into the plant cytoplasm (Figure S5). Treatment with the cell death inducer Npp1 alone was sufficient to trigger multiple cellular stress responses, including autophagy, senescence, apoptosis-like processes, and DDR activation. This supports the idea that all PCD pathways in plants may be interconnected. As in vertebrates, invertebrate MIFs are attributed a wide range of functions, including roles in cross-kingdom cell death, immunity, homeostasis, development, cell proliferation and wound healing. However, the molecular pathways underlying these functions remain poorly characterized in non-mammalian organisms. This raises an intriguing question of how MIF proteins exert similar biological roles across vastly different organisms, especially given that most known mammalian MIF receptors are not conserved in other phyla. Our study provides the first evidence of a direct interaction between an insect MIF and the plant transcription factor SOG1, comparable to MIF and p53 interaction in mammals. Despite the lack of a sequence similarity between SOG1 and p53 [49], MpMIF1 ability to bind SOG1 highlights both functional convergence and evolutionary questions. These findings emphasize MIF as a remarkably versatile and evolutionary intriguing molecule [9,12], capable of modulating fundamental stress-response pathways across biological kingdoms. Together, these findings highlight MpMIF1 not only as a suppressor of aphid-triggered plant cell death but also as a potential biotechnological tool to manipulate DDR–SOG1–PCD signaling in crops, ultimately offering novel strategies for enhancing plant resilience against aphid infestation.

**Figure 7:**
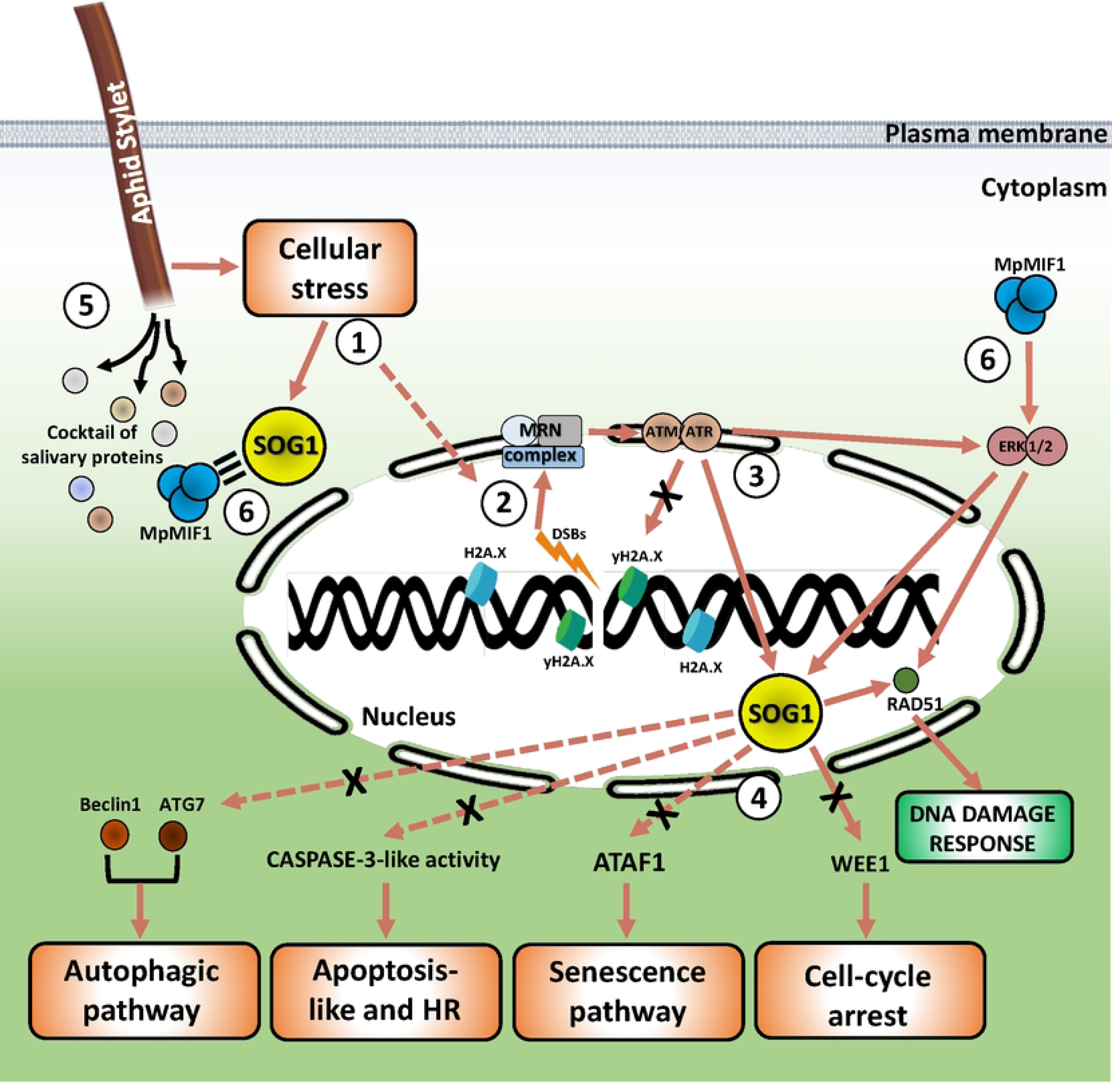
Model of PCD and DDR signaling pathways affected by MIF. This schematic model shows the candidate genes analyzed in our study and their interactions. Documented interactions are shown as plain arrows while likely interactions are shown as dashed arrows. The effect of MpMIF1 based on our results, are shown as black crosses. (1) Stylet punctures cause significant cellular stress (Figure S5), leading to the activation of SOG1 and potentially inducing DNA damage thus activating the entire network responsible for the detection and repair of double-strand break. (2) In the presence of double-strand breaks (DSBs), the MRN complex acts as the primary sensor, facilitating the recruitment and activation of the kinases ATM and ATR. These kinases phosphorylate γH2AX, which then acts as a scaffold for the recruitment of DNA repair proteins, including RAD51. (3) ATM and ATR also phosphorylate and activate multiple downstream effectors involved in the DNA damage response. Notably, they target SOG1. This phosphorylation event is further enhanced by the activation of MAPKs ERK1/2, which are also downstream of ATM/ATR signaling. (4) Activation (translocation) of SOG1 induces: autophagic responses via activation of ATG7 and BECLIN1; apoptosis-like pathways through DEVDase (caspase-3-like) activity; senescence programs involving ATAF1; inhibition of cell cycle progression via WEE1 kinase, and activates or inhibits DNA repair by regulating RAD51. (5) Shortly after penetration, the aphid stylet secretes a cocktail of salivary proteins, containing MpMIF1. (6) MpMIF1 physically interacts with SOG1. This interaction likely prevents translocation of SOG1 in the nucleus and the subsequent activation of the downstream pathways, thereby inhibiting its pro-death activity and indirectly promoting DNA damage repair mechanisms. MpMIF1 also sustains MAPK ERK 1/2 phosphorylation that could increase RAD51 stability and participates indirectly to DDR progression.

## Acknowledgments

We thank Oliver Pierre and Julie Soltys for assistance with microscopy and Sophie Pagnota with TEM analysis (CCMA, “Université Côte D’Azur”). We are grateful to technical help of Roxane Pichot and Oliver Migliore for biochemistry and plant culture, respectively.

## Author Contribution

KM, CC and JdAE conceived the project and designed the experiments. CAL, KM and JdAE performed the microscopic experiments. AF helped with RT-qPCR, TF for Gateway cloning and GDS with Western Blot optimization. KM performed remaining experiments. KM, CC and JdAE analyzed the data, KM wrote the manuscript and CC and JdAE amended the paper.

## Data availability

The data that supports the findings of this study are available upon request.

## Supporting Information

***Table S1:*** Primers used in qPCR.

***Table S2:*** Primers used for TAGs line creation using gateway cloning technology.

***Figure S1: Subcellular localization of MpMIF1 following agroinfiltration in plant cells.***

Confocal microscopic pictures showing subcellular localization of MpMIF1: GFP line following agroinfiltration in plant cells at two days post infiltration. *Myzus persicae* MIF1 (*MpMIF1* GenBank: KP218519.1) sequences were cloned into the entry vector pDON207 and transferred into the destination vectors pK7WGF2.0 already containing GFP sequence. Using the same agroinfiltration protocol described in the Materials and Methods section, we successfully expressed the MpMIF1::GFP fusion protein in plant tissues. Images were acquired with a Zeiss LSM780 confocal microscope equipped with a 20× Plan-Apochromat objective. GFP fluorescence was excited with an argon laser at 488 nm, and emission was collected between 493 and 555 nm. The resulting images show that, following agroinfiltration, MpMIF1 localizes predominantly in the cytoplasm and the nuclei (white arrows). Scale bars = 70 μm.

***Figure S2: Effect of caspase-3 inhibitor AC-DEVD-CHO on caspase-3-like activities in plants.***

Caspase-3-like activity at 15, 20, and 24 hours’ post-infiltration (HPI), with the presence of caspase-3 inhibitor AC-DEVD-CHO (lighter color on the right side of each normal activity bar). The inhibitor addition experiment was performed under the same conditions as the standard caspase-3-like activity assay (see Materials and Methods), with 50 µM AC-DEVD-CHO added to each well. No significant statistical differences were observed between samples with or without the inhibitor.

***Figure S3: MIF-TAG cell death inhibition conservation.***

(A) Leaves images showing cell death in *N. benthamiana*. Leaves were inoculated with different strains of *A. tumefaciens* harboring *MpMIF1-HA* or *MpMIF1-CMyc* construct combined with *Npp1*. For leaves pictures, the visible symptoms of Npp1-induced cell death are dehydration and brown lesions. These symptoms do not establish in leaf areas that express MpMIF1 or MpMIF1-TAGs showing a conservation of function. Pictures were taken at 1 and 3-Days Post infiltration (DPI) with *A. tumefaciens*. Bars = 200 μm (B) Chlorophyll quantification in plant leaves after agroinfiltration. The total chlorophyll was measured at 1 and 3 DPI in *N. benthamiana* plant leaves subjected to the same treatment as above. Error bars represent the Standard Error of 12 biological replicates. The asterisks indicate statistical differences according to ANOVA and Tukey-Kramer test (***p < 0.0001, **p < 0.01, *p < 0.05, ns = not significant).

***Figure S4: Split-luciferase assay between MpMIF1-nLUC; cLUC-ATAF1 and cLUC-SOG1.***

MpMIF1-nLUC was transiently co-expressed at 1 and 2 DPI with cLUC-ATAF1 and cLUC-SOG1. (A) Black spots correspond to detected luminescence signal under CCD-camera to show luciferase activities. (B) Box plot showing luciferase activity from the split-luciferase complementation assay using Luminometer. The luminescence of seven independent experiments was quantified in single 4 mm punch disks placed into 96 well plate and incubated with 1mM luciferin. For all diagrams, error bars represent the Standard Error and the asterisks indicate statistical differences according to ANOVA and Tukey-Kramer test (***p < 0.0001, **p < 0.01, *p < 0.05, ns = not significant).

***Figure S5: In vivo-Aphids Myzus Persicae infection affects different cell death pathways.***

Our study, using the heterologous expression of MpMIF1 showed that the expression of MpMIF1 influences several plant cell death and DNA damage response pathways. To establish a connection between the effects observed under heterologous expression and in vivo aphid (Myzus persicae) infection, we investigated the impact of aphid infection on the expression of four of the plant cell death-related candidates: *ATG7, BECLIN1, ATAF1,* and *SOG1*. (A) Schematic representation of the experimental design. M. persicae were reared on N. benthamiana under long photoperiod (16 hours’ light/ 8 hours’ dark) and elevated temperature conditions (19°C) to maintain continuous parthenogenesis. Only parthenogenetic apterous adults were used for all experiments. Between fifty and seventy individuals were placed under clip cages on N. benthamiana plants. After one or two days of infection, aphids were removed and plant discs were collected following a timing kinetics: 0min, 5mn; 4hours; 24hours and 48hours after removal of aphids. As control, a non-infected treatment was performed using an empty clip cage maintained for 1 or 2 days to measure effect of clip-cage mechanical stress. Total RNA from N. benthamiana tissues was then extracted and expression of plant specific cell death genes were analyzed using RT-QPCR: *ATG7, BECLIN1 SOG1* and *ATAF1*. mRNA expression ratios were calculated relative to references genes for each sample: *Ef1a* (Elongation Factor 1), *UBQr* (Ribosomal Ubiquitin) and *GAPDH* (Glyceraldehyde 3-phosphate dehydrogenase). Six biological replicates were performed and data were submitted to statistical analysis using GraphPad prism. (B) Expression ratios of *ATG7, BECLIN1 SOG1* and *ATAF1* in tissues of plant infected for 1 day (blue) or two days (green). Six biological replicates were performed and data were submitted to statistical analysis using GraphPad prism. Values marked by different letters are significantly different (p < 0.001; ANOVA and Tukey-Kramer test). Values marked by identical letters are not significantly different.

For the autophagy-related genes *ATG7* and *BECLIN1*, no significant differences were observed after one day of aphid infection. In samples exposed to infection for 2 days, *ATG7* expression decreased shortly after aphid removal and returned to basal levels 24 hours after aphid removal. Interestingly, *BECLIN1* showed a different pattern, with a continuous increase in expression over time, peaking 48 hours after aphid removal, in samples infected for 2 days.

Regarding the senescence-related gene *ATAF1*, no significant differences were observed in samples infected during one day. In contrast, plant tissues exposed to aphids for 2 days showed an increased expression at 24 hours after aphid removal, before returning to basal levels at 48 hours. Aphid infection for one day also led to a significant increase in *SOG1* expression 24 hours after aphid removal. In the 2-days infection condition, *SOG1* expression decreased immediately after aphid removal but rapidly returned to basal levels. These RT-qPCR results indicate that aphid infection affects the expression of multiple plant cell death genes involved in different pathways (autophagy and senescence). The increase in cell death gene expression after aphid removal suggests that aphid forced detachment causes cellular damage. Conversely, the lower expression of cell death genes when aphids were still present (T0 min) highlights the ability of aphids to suppress plant immune responses—likely through the secretion of salivary proteins such as MpMIF1. Particular attention was given to *SOG1*, given its role as a functional analog of p53 in plants. Our results confirm that *in vivo* aphid infection also affects *SOG1* expression, further supporting its involvement in the plant’s DNA damage response against piercing and sucking parasite.

## References

1. Wood CL, Johnson PT. A world without parasites: exploring the hidden ecology of infection. Frontiers in Ecology and the Environment 13 (2015): 425–434.

2. Xu Y, Gray SM. Aphids and their transmitted potato viruses: A continuous challenges in potato crops. Journal of Integrative Agriculture 19 (2020): 367–375.

3. Giordanengo PP, Brunissen L, Rusterucci C, et al. Compatible plant-aphid interactions: How aphids manipulate plant responses. Comptes Rendus. Biologies 333 (2010): 516–523.

4. Mutti NS, Louis J, Pappan LK, et al. A protein from the salivary glands of the pea aphid, *Acyrthosiphon pisum* is essential in feeding on a host plant. Proc Natl Acad Sci USA 105 (2008): 9965–9969.

5. Guerrieri E, Digilio MC. Aphid-plant interactions: a review. J of Plant Interactions 3 (2008): 223–232.

6. Saheed SA, Liu L, Jonsson L, et al. Xylem – as well as phloem – sustains severe damage due to feeding by the Russian wheat aphid. South African Journal of Botany 73 (2007): 593–599.

7. Morkunas I, Mai VC, Gabryś B. Phytohormonal signaling in plant responses to aphid feeding. Acta Physiol Plant 33 (2011): 2057–2073.

8. Naessens E, Dubreuil G, Giordanengo P, et al. A Secreted MIF Cytokine Enables Aphid Feeding and Represses Plant Immune Responses. Current Biology 25 (2015): 1898–1903.

9. Sparkes A, De Baetselier P, Roelants K, et al. Reprint of: The non-mammalian MIF superfamily. Immunobiology 222 (2017): 858–867.

10. Michelet C, Danchin EGJ, Jaouannet M, et al. Cross-Kingdom Analysis of Diversity, Evolutionary History, and Site Selection within the Eukaryotic Macrophage Migration Inhibitory Factor Superfamily. Genes 10 (2019): 740.

11. Gruner K, Leissing F, Sinitski D, et al. Chemokine-like MDL proteins modulate flowering time and innate immunity in plants. Journal of Biological Chemistry 296 (2021): 100611.

12. Bucala R. A most interesting factor. Nature 408 (2000): 146–147.

13. Calandra T, Roger T. Macrophage migration inhibitory factor: a regulator of innate immunity. Nat Rev Immunol 3 (2003): 791–800.

14. Huang S, Cao Y, Lu M, et al. Identification and functional characterization of *Oncomelania hupensis* macrophage migration inhibitory factor involved in the snail host innate immune response to the parasite *Schistosoma japonicum*. International Journal for Parasitology 47 (2017): 485–499.

15. Cho Y, Jones BF, Vermeire JJ, et al. Structural and Functional Characterization of a Secreted Hookworm Macrophage Migration Inhibitory Factor (MIF) That Interacts with the Human MIF Receptor CD74. Journal of Biological Chemistry 282 (2007): 23447–23456.

16. Zhao J, Li L, Liu Q, et al. A MIF-like effector suppresses plant immunity and facilitates nematode parasitism by interacting with plant annexins. Journal of Experimental Botany 70 (2019): 5943–5958.

17. Holowka T, Castilho TM, Garcia AB, et al. *Leishmania* -encoded orthologs of macrophage migration inhibitory factor regulate host immunity to promote parasite persistence. FASEB J 30 (2016): 2249–2265.

18. Jaworski DC, Jasinskas A, Metz CN, et al. Identification and characterization of a homologue of the pro-inflammatory cytokine Macrophage Migration Inhibitory Factor in the tick, *Amblyomma americanum*. Insect Molecular Biology 10 (2001): 323–331.

19. Dubreuil G, Deleury E, Crochard D, et al. Diversification of MIF immune regulators in aphids: link with agonistic and antagonistic interactions. BMC Genomics 15 (2014): 762.

20. Reymond P, Calandra T. Plant Immune Responses: Aphids Strike Back. Current Biology 25 (2015): R604–R606.

21. Mitchell RA, Liao H, Chesney J, et al. Macrophage migration inhibitory factor (MIF) sustains macrophage proinflammatory function by inhibiting p53: Regulatory role in the innate immune response. Proc Natl Acad Sci USA 99 (2002): 345–350.

22. Hudson JD, Shoaibi MA, Maestro R, et al. A Proinflammatory Cytokine Inhibits P53 Tumor Suppressor Activity. The Journal of Experimental Medicine 190 (1999): 1375–1382.

23. Jung H, Seong HA, Ha H. Critical Role of Cysteine Residue 81 of Macrophage Migration Inhibitory Factor (MIF) in MIF-induced Inhibition of p53 Activity. Journal of Biological Chemistry 283 (2008): 20383–20396.

24. Brock SE, Rendon BE, Xin D, et al. MIF Family Members Cooperatively Inhibit p53 Expression and Activity. PLoS ONE 9 (2014): e99795.

25. Lue H, Kleemann R, Calandra T, et al. Macrophage migration inhibitory factor (MIF): mechanisms of action and role in disease. Microbes and Infection 4 (2002): 449–460.

26. Xu L, Li Y, Sun H, et al. Current developments of macrophage migration inhibitory factor (MIF) inhibitors. Drug Discovery Today 18 (2013): 592–600.

27. Lam, E. Controlled cell death, plant survival and development. Nat Rev Mol Cell Biol 5 (2004): 305–315.

28. Reape TJ, Molony EM, McCabe PF. Programmed cell death in plants: distinguishing between different modes. J Exp Bot 59 (2008): 435–444.

29. Kroemer G, Galluzzi L, Vandenabeele P, et al. Classification of cell death: recommendations of the Nomenclature Committee on Cell Death 2009. Cell Death and Differentiation 16 (2009): 3–11.

30. Mukhtar MS, McCormack ME, Argueso CT, et al. Pathogen Tactics to Manipulate Plant Cell Death. Current Biology 26 (2016): R608–R619.

31. Emanuele S, Oddo E, D’Anneo A, et al. Routes to cell death in animal and plant kingdoms: from classic apoptosis to alternative ways to die—a review. Rend Fis Acc Lincei 29 (2018): 397409.

32. Bozhkov P, Jansson C. Autophagy and Cell-Death Proteases in Plants: Two Wheels of a Funeral Cart. Autophagy 3 (2007): 136–138.

33. Xu Q, Zhang L. Plant caspase-like proteases in plant programmed cell death. Plant Signaling and Behavior 4 (2009): 902–904.

34. Iakimova ET, Michalczuk L, Woltering EJ. Hypersensitive cell death in plants – its mechanisms and role in plant defence against pathogens 13 (2005).

35. Coll NS, Epple P, Dangl JL. Programmed cell death in the plant immune system. Cell Death Differ 18 (2011): 1247–1256.

36. Roudaire T, Héloir MC, Wendehenne D, et al. Cross Kingdom Immunity: The Role of Immune Receptors and Downstream Signaling in Animal and Plant Cell Death. Frontiers in Immunology 11 (2021): 612452.

37. Zheng W, Xie W, Yin D, et al. ATG5 and ATG7 induced autophagy interplays with UPR via PERK signaling. Journal of Cell Communication and Signaling 17 (2019): 42.

38. Tran S, Fairlie WD, Lee EF. BECLIN1: Protein Structure, Function and Regulation. Cells 10 (2021): 1522.

39. Van Doorn WG, Beers EP, Dangl JL, et al. Morphological classification of plant cell deaths. Cell Death and Differentiation 18 (2011): 1241–1246.

40. Minina EA, Dauphinee AN, Ballhaus F, et al. Apoptosis is not conserved in plants as revealed by critical examination of a model for plant apoptosis-like cell death. BMC Biol 19 (2021): 100.

41. Dickman M, Williams, B, Li Y, et al. Reassessing apoptosis in plants. Nature Plants 3 (2017): 773–779.

42. Meschichi A, Zhao L, Reeck S, et al. The plant-specific DDR factor SOG1 increases chromatin mobility in response to DNA damage. EMBO Reports 23 (2022): e54736.

43. Mahapatra K, Roy S. An insight into the mechanism of DNA damage response in plants-role of Suppressor Of Gamma Response 1: An overview. Mutation Research/Fundamental and Molecular Mechanisms of Mutagenesis 819–820 (2020): 111689.

44. Lavin MF, Delia D, Chessa L. ATM and the DNA damage response: Workshop on Ataxia-Telangiectasia and Related Syndromes. EMBO Reports 7 (2006): 154–160.

45. Burma S, Chen BP, Murphy M, et al. ATM Phosphorylates Histone H2AX in Response to DNA Double-strand Breaks. Journal of Biological Chemistry 276 (2001): 42462–42467.

46. Rogakou EP, Pilch DR, Orr AH, et al. DNA Double-stranded Breaks Induce Histone H2AX Phosphorylation on Serine 139. Journal of Biological Chemistry 273 (1998): 5858–5868.

47. Tashiro S, Walter J, Shinohara A, et al. Rad51 Accumulation at Sites of DNA Damage and in Postreplicative Chromatin. Journal of Cell Biology 150 (2000): 283–292.

48. Liu F, Xu Y, Zhou L, et al. DNA Repair Gene ZmRAD51A Improves Rice and Arabidopsis Resistance to Disease. IJMS 20 (2019): 807.

49. Yoshiyama K, Sakaguchi K, Kimura S. DNA Damage Response in Plants: Conserved and Variable Response Compared to Animals. Biology 2 (2013): 1338–1356.

50. Hu Z, Cools T, De Veylder L. Mechanisms Used by Plants to Cope with DNA Damage. Annu Rev Plant Biol 67 (2016): 439–62.

51. Yoshiyama KO, Kimura S, Maki H, et al. The role of SOG1, a plant-specific transcriptional regulator, in the DNA damage response. Plant Signaling and Behavior 9 (2014): e28889.

52. Yoshiyama KO. SOG1: a master regulator of the DNA damage response in plants. Genes and Genetic Systems 90 (2015): 209–216.

53. Avila-Ospina L, Moison M, Yoshimoto K, Masclaux-Daubresse C. Autophagy, plant senescence, and nutrient recycling. Journal of Experimental Botany 65 (2014): 3799–3811.

54. Garapati P, Xue GP, Munné-Bosch S, et al. Transcription Factor ATAF1 in *Arabidopsis* Promotes Senescence by Direct Regulation of Key Chloroplast Maintenance and Senescence Transcriptional Cascades. Plant Physiol 168 (2015): 1122–1139.

55. Cabral D, Banora MY, Antonino JD, et al. The plant WEE1 kinase is involved in checkpoint control activation in nematode-induced galls. New Phytologist 225 (2020): 430–447.

56. Wang L, Zhan L, Zhao Y, et al. The ATR-WEE1 kinase module inhibits the MAC complex to regulate replication stress response. Nucleic Acids Research 49 (2021): 1411–1425.

57. Oome S, Raaymakers TM, Cabral A, et al. Nep1-like proteins from three kingdoms of life act as a microbe-associated molecular pattern in *Arabidopsis*. Proc Natl Acad Sci USA 111 (2014): 16955–16960.

58. Lenarčič T, Pirc K, Hodnik V, et al. Molecular basis for functional diversity among microbial Nep1-like proteins. PLoS Pathog 15 (2019): e1007951.

59. Jaouannet M, Pavaux AS, Pagnotta S, et al. Atypical Membrane-Anchored Cytokine MIF in a Marine Dinoflagellate. Microorganisms 8 (2020): 1263.

60. De Almeida Engler J, Van Poucke K, Karimi M, et al. Dynamic cytoskeleton rearrangements in giant cells and syncytia of nematode-infected roots. The Plant Journal 38 (2004): 12–26.

61. Yoshiyama KO, Kobayashi J, Ogita N, et al. ATM-mediated phosphorylation of SOG1 is essential for the DNA damage response in *Arabidopsis*. EMBO Reports 14 (2013): 817–822.

62. Otegui MS, Noh Y, Martínez DE, et al. Senescence-associated vacuoles with intense proteolytic activity develop in leaves of *Arabidopsis* and soybean. The Plant Journal 41 (2005): 831–844.

63. Tran V, Weier D, Radchuk R, et al. Caspase-Like Activities Accompany Programmed Cell Death Events in Developing Barley Grains. PLoS ONE 9 (2014): e109426.

64. Flott S, Kwon Y, Pigli YZ, et al. Regulation of Rad51 function by phosphorylation. EMBO Reports 12 (2011): 833–839.

65. De Schutter K, Joubès J, Cools T, et al. *Arabidopsis* WEE1 Kinase Controls Cell Cycle Arrest in Response to Activation of the DNA Integrity Checkpoint. The Plant Cell 19 (2007): 211–225.

66. Mugford ST, Barclay E, Drurey C, et al. An Immuno-Suppressive Aphid Saliva Protein Is Delivered into the Cytosol of Plant Mesophyll Cells During Feeding. MPMI 29 (2016): 854–861.

67. Zechmann B. Ultrastructure of plastids serves as reliable abiotic and biotic stress marker. PLoS ONE 14 (2019): e0214811.

68. Zavafer A, González-Solís A, Palacios-Bahena S, et al. Organized Disassembly of Photosynthesis During Programmed Cell Death Mediated By Long Chain Bases. Scientific Reports 10 (2020): 10360.

69. Howell SH. Endoplasmic Reticulum Stress Responses in Plants. Annu. Rev. Plant Biol 64 (2013): 477–499.

70. Qiang X, Zechmann B, Reitz MU, et al. The Mutualistic Fungus *Piriformospora indica* Colonizes *Arabidopsis* Roots by Inducing an Endoplasmic Reticulum Stress–Triggered Caspase-Dependent Cell Death. Plant Cell 24 (2012): 794–809.

71. Smertenko A, Franklin-Tong VE. Organisation and regulation of the cytoskeleton in plant programmed cell death. Cell Death and Differentiation 18 (2011): 1263–1270.

72. Manova V, Gruszka D. DNA damage and repair in plants – from models to crops. Front. Plant Sci 6 (2015).

73. Liu Y, Schiff M, Czymmek K, et al. Autophagy Regulates Programmed Cell Death during the Plant Innate Immune Response. Cell 121 (2005): 567–577.

74. Li F, Zhang C, Li Y, et al. Beclin1 restricts RNA virus infection in plants through suppression and degradation of the viral polymerase. Nat Commun 9 (2018): 1268.

75. Lang T, Foote A, Lee JPW, et al. MIF: Implications in the Pathoetiology of Systemic Lupus Erythematosus. Front Immunol 6 (2015).

76. Del Duca S, Serafini-Fracassini D, Cai G. Senescence and programmed cell death in plants: polyamine action mediated by transglutaminase. Front. Plant Sci 5 (2014).

77. Sauler M, Bucala R, Lee PJ. Role of macrophage migration inhibitory factor in age-related lung disease. American Journal of Physiology-Lung Cellular and Molecular Physiology 309 (2015): L1–L10.

78. Paluch-Lubawa E, Stolarska E, Sobieszczuk-Nowicka E. Dark-Induced Barley Leaf Senescence – A Crop System for Studying Senescence and Autophagy Mechanisms. Front Plant Sci 12 (2021): 635619.

79. Bonneau L, Ge Y, Drury GE, et al. What happened to plant caspases? J Exp Bot 59 (2008): 491499.

80. Chen D, Zhao C, Zhang J, et al. Small Molecule MIF Modulation Enhances Ferroptosis by Impairing DNA Repair Mechanisms. AdVanced Science 11 (2024): 2403963.

81. Subbannayya T, Variar P, Advani J, et al. An integrated signal transduction network of macrophage migration inhibitory factor. Journal of Cell Communication and Signaling 10 (2016): 165–170.

82. Taj G, Agarwal P, Grant M, et al. MAPK machinery in plants: Recognition and response to different stresses through multiple signal transduction pathways. Plant Signaling and Behavior 5 (2010): 1370–1378.

83. Meng X, Zhang S. MAPK Cascades in Plant Disease Resistance Signaling. Annu. Rev. Phytopathol 51 (2013): 245–266.

84. Yue J, López JM. Understanding MAPK Signaling Pathways in Apoptosis. International Journal of Molecular Sciences 21 (2020): 2346.

85. Allan LA, Morrice N, Brady S, et al. Inhibition of caspase-9 through phosphorylation at Thr 125 by ERK MAPK. Nat Cell Biol 5 (2003): 647–654.

86. Su YJ, Tsai MS, Kuo YH, et al. Role of Rad51 Down-Regulation and Extracellular Signal-Regulated Kinases 1 and 2 Inactivation in Emodin and Mitomycin C-Induced Synergistic Cytotoxicity in Human Non-Small-Cell Lung Cancer Cells. Molecular Pharmacology 77 (2010): 633–643.

87. Wu GS. The Functional Interactions Between the MAPK and P53 Signaling Pathways. Cancer Biology and Therapy 3 (2004): 156–161.

88. Sinitski D, Gruner K, Brandhofer M, et al. Cross-kingdom mimicry of the receptor signaling and leukocyte recruitment activity of a human cytokine by its plant orthologs. Journal of Biological Chemistry 295 (2020): 850–867.

89. Horn HF, Vousden KH. Coping with stress: multiple ways to activate p53. Oncogene 26 (2007): 1306–1316.

90. Baeza Garcia A, Pierce RJ, Gourbal B, et al. Involvement of the Cytokine MIF in the Snail Host Immune Response to the Parasite *Schistosoma mansoni*. PLoS Pathog 6 (2010): e1001115.

